# Single-nucleus RNA sequencing provides insights into the GL261-GSC syngeneic mouse model of glioblastoma

**DOI:** 10.1101/2023.10.26.564166

**Authors:** Laura García-Vicente, Michael Borja, Vanessa Tran, Andrea Álvarez-Vázquez, Raquel Flores-Hernández, Alejandro Granados, Aaron McGeever, Yang-Joon Kim, Leah Dorman, Angela Detweiler, Honey Mekonen, Sheryl Paul, Angela O Pisco, Norma F Neff, Arantxa Tabernero

## Abstract

Glioblastoma (GBM) is an aggressive tumor with very bad prognosis. The urgent need to find new effective therapies is challenged by the unique characteristics of GBM, including high intra and intertumoral heterogeneity. Using single-nucleus transcriptomics (snRNA-seq), we characterized the panorama of a preclinical immunocompetent murine model based in the implantation of mouse glioblastoma stem cells (GL261-GSCs) into the brain parenchyma. Additionally, we performed Visium spatial transcriptomics in the in vivo model to confirm the location of annotated cells. To understand the technical bias of this approach, we performed two scRNA-seq methods in GBM cells. We thoroughly characterized the tumor microenvironment (TME) at early and late stages of tumor development and upon treatment with temozolomide (TMZ), the standard of care for patients with GBM, and with Tat-Cx43_266-283_, a promising experimental treatment. We identified prominent GBM targets that can be addressed using this preclinical model, such as *Grik2*, *Nlgn3*, *Gap43* or *Kcnn4*, which are involved in electrical and synaptic integration of GBM cells into neural circuits, as well as the expression of *Nt5e*, *Cd274* or *Irf8*, which indicates the development of immune evasive properties in these GBM cells. In agreement, snRNA-seq unveiled high expression of several immunosuppressive-associated molecules in immune cells, such as *Csf1r, Arg1, Mrc1* and *Tgfb1*, suggesting the development of an immunosuppressive microenvironment. We also show the landscape of cytokines, cytokine receptors, checkpoint ligands and receptors in tumor and TME cells, which are crucial data for a rational design of immunotherapy studies. Thus, Mrc1, PD-L1, TIM-3 or B7-H3 are among the immunotherapy targets that can be addressed in this model. Finally, the comparison of the preclinical GL261-GSC GBM model with human GBM subtypes unveiled important similarities with the recently identified TMEmed human GBM, indicating that preclinical data obtained in GL261-GSC GBM model might be applied to TMEmed human GBM, improving patient stratification in clinical trials. In conclusion, this work provides crucial information for future preclinical studies in GBM improving their clinical application.

## Introduction

GBM is the most aggressive and frequent form of primary brain tumor and, despite continuous effort to find an effective treatment, is considered as one of the deadliest types of cancer, with a median survival of only 16 months. GBMs are characterized by fast and aggressive growth, high infiltrative capacity, and resistance to current treatments. The main challenges underlying therapeutic failure are derived from its cellular and molecular heterogeneity. Fortunately, increasing knowledge of cellular and molecular alterations in brain tumors has improved their classification ^1^ with benefits in personalized treatments. For instance, inhibitors of mutant IDH1 and IDH2 enzymes have shown positive results in clinical trials for IDH-mutant gliomas ^2^. Regarding IDHwt GBM, recent studies have identified three novel subtypes with significantly different tumor microenvironment (TME) compositions and different response to immunotherapy treatments ^3^, which supports that deciphering GBM heterogeneity may improve their treatment. High-throughput studies, such as single-cell RNA sequencing (scRNA-Seq) studies have helped to clarify intra-tumor and inter-tumor heterogeneity ^4–6^. scRNA-Seq has emerged as a key method to characterize cellular states and is continuously used to analyze tumor samples ^7^; however, it requires a quick dissociation of fresh tissue and enzymatic digestion that can damage sensitive cell types such as neurons ^8^. Conversely, single-nucleus RNA-Seq (snRNA-Seq) avoids the aggressive enzymatic digestion, preserving the information from delicate cell types ^9, 10^, such as neurons in TME.

Unfortunately, many experimental treatments that were successful in preclinical models have failed in subsequent clinical trials ^11^. Among the causes for this failure are the differences between preclinical models and human glioblastoma. A profound cellular and molecular characterization of GBM preclinical models and the correlation with human GBM subtypes is required to translate the preclinical results to the most similar GBM subtype. The GL261 model is the most widely used syngeneic model of glioblastoma due to a series of advantages, such as the possibility to study the immune system. It has been widely used in studies of gene therapy, immune cell transfer, monoclonal antibodies, cytokine therapies, checkpoint inhibitors, and dendritic vaccines ^12–16^. Intracranially implanted GL261 cells grow rapidly and form tumors with 100% penetrance and, although classified as GBM because of its aggressivity, it displays few histopathological GBM indicators ^17^. GBMs contain populations of cells with stem cell properties, known as GBM stem cells (GSCs), which are highly tumorigenic, possess tumor-propagating potential, and exhibit resistance to chemotherapy and radiotherapy ^18, 19^. Yi and colleagues described a GBM model in which the tumor is initiated by the implantation of GSCs obtained from GL261 cells in immunocompetent mice (GL261-GSC GBM model), which recapitulates the typical features of human GBM ^20, 21^.

Here, we use snRNA-Seq to create a map of murine GBM cellular states from in vitro to in vivo using the GL261-GSC GBM model. We performed snRNA-Seq in a total of 14 samples from tumor-bearing mice and 5 samples from cultured GBM cells, using a microfluidic-droplet-based method. Additionally, we performed Visium spatial transcriptomics in the in vivo model to confirm the location of annotated cells. To understand the technical bias of this approach, we performed two scRNA-seq methods in GBM cells. We thoroughly characterized the TME in the GL261-GSC GBM model at early and late stages of GBM development and upon treatment with temozolomide (TMZ), the actual standard of care chemotherapeutic drug for GBM, and a novel experimental treatment, the cell penetrating peptide Tat-Cx43_266-283_ ^21–24^. The present study identified prominent GBM targets that can be addressed using this preclinical model. Furthermore, we unveiled the similarities of GL261-GSC GBM model with the most frequent human GBM subtype, which might improve the translation of preclinical results to a more specific clinical setting. In conclusion, this work provides crucial information for future preclinical studies in GBM improving their clinical application.

## Results

### Identification of tumor and non-tumor cells in the GL261-GSC GBM model

To characterize the biology of murine GBM cells growing in the brain TME, 5,000 GL261-GSCs were intracranially injected into the brains of syngeneic C57BL/6 mice. This low number of cells develops very aggressive tumors with poor mice survival, indicating high malignancy of GL261-GSCs ^21^. We collected samples of four GBM-bearing mouse brains, including tumor core and surrounding tissue, at two stages of tumor growth, early (7 days post-implantation) and late (28 days post-implantation), and isolated nuclei for snRNA-Seq. This process is described in detail in the Methods section and graphically summarized in Figure 1.

**Figure 1.**
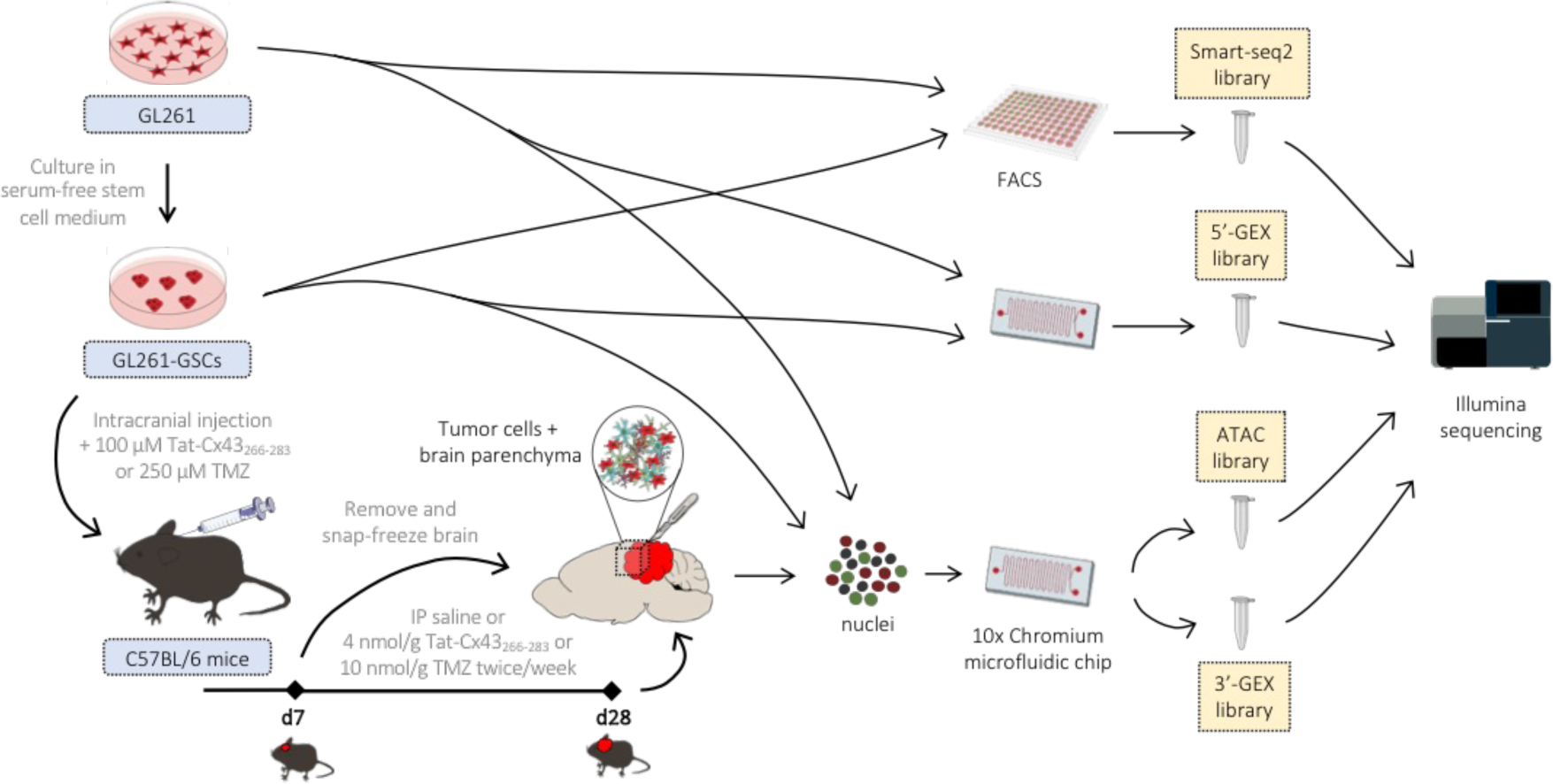
Study overview. GL261-GSCs were grown as neurospheres in stem cell medium. They were obtained from GL261 cells grown in adherence in differentiation medium. 5,000 GL261-GSCs were intracranially injected into the brains of syngeneic C57BL/6 mice in saline solution or 100 µM Tat-Cx43_266-283_ or 250 µM temozolomide (TMZ). After 7 days, these treatments were intraperitoneally (IP) administered twice per week. Samples were collected from GBM-bearing mouse brains, including tumor core and surrounding tissue, at two stages of tumor growth, early (7 days post-implantation) and late (28 days post-implantation), and isolated nuclei for snRNA-Seq. scRNA-seq was performed in GL261 and GL261-GSCs while snRNA-Seq was performed in all the in vitro and in vivo samples.

At 7 days post-implantation, tumors are small, and we were able to take samples containing the whole tumor and its TME. In the case of 28-days tumors, we obtained samples from the border of the tumor to capture tumor cells as well as the other cell types present in the brain parenchyma. Additionally, we analyzed 2 samples of GL261 cells growing in culture in differentiation conditions, and 3 samples of GL261-GSCs growing as neurospheres in stem cell medium, the condition in which they grew prior to implantation into the brain. In total, 28,833 cells passed our quality controls and were further analyzed. We used uniform manifold approximation and projection (UMAP) for dimensionality reduction ^25^, followed by Leiden clustering ^26^, which resulted in the identification of 22 separate clusters (Figure 2A). We used hierarchical clustering and a Wilcoxon rank sum test to obtain differentially expressed genes (DEGs) in each cluster (Figure 2B and Table S1). Clusters were classified by high expression of gene markers from published mouse CNS cell type databases ^27^ (Figure 2C). All clusters showed high expression of characteristic gene sets except for those corresponding to cultured GL261 cells (clusters 0, 1, 2, 4, 5, 6, 11, and 19), GL261-GSCs (clusters 1, 2, 6, and 11), and cluster 0, which we preliminarily classified as malignant (implanted GL261-GSCs).

**Figure 2.**
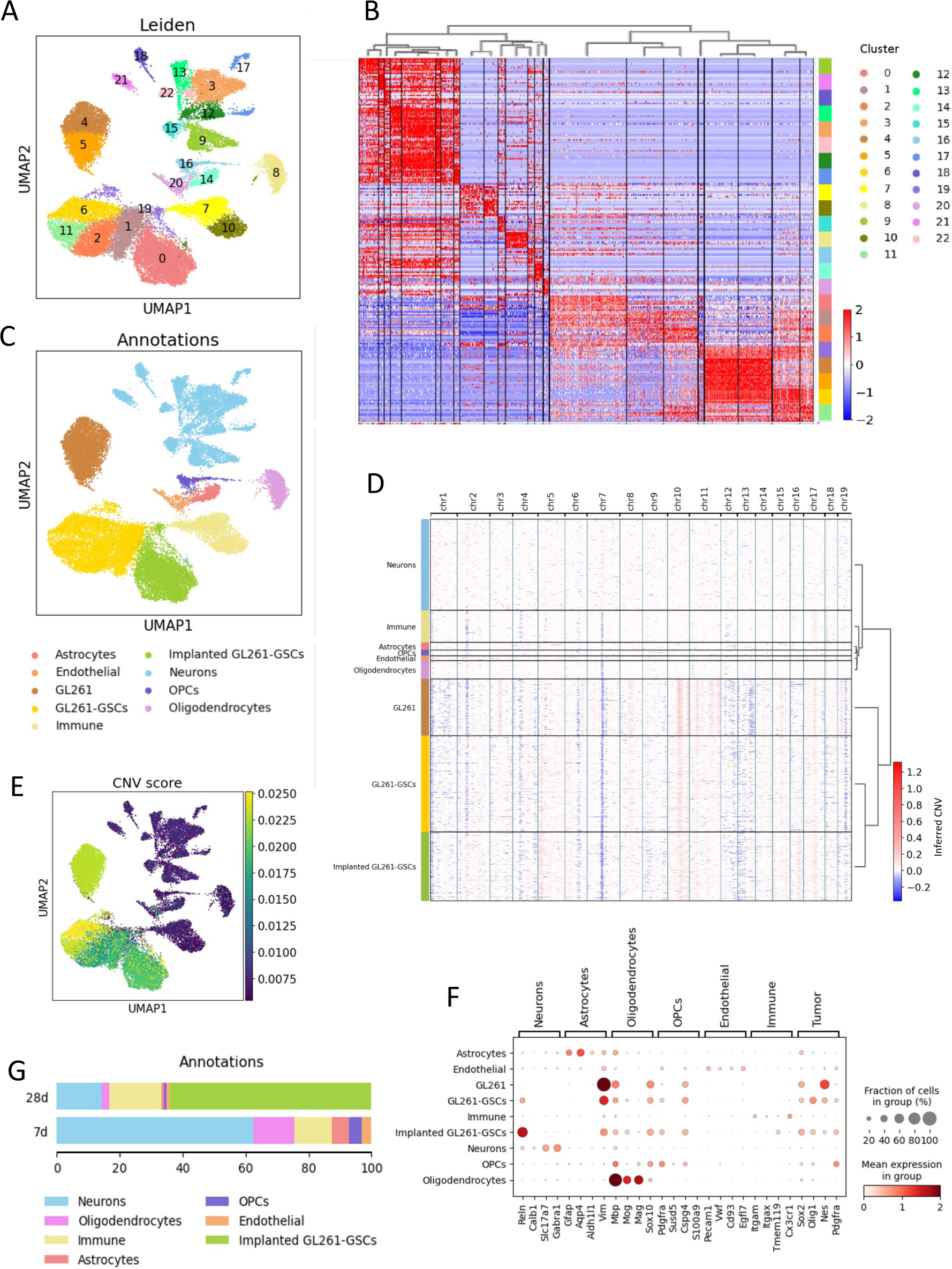
snRNA-Seq of GL261-GSCs and tumor samples. (A) UMAP visualization of all cells collected, colored by Leiden clustering. k = 28,833 individual cells (B) Heatmap showing relative expression of each of the 10 top expressed genes in each Leiden cluster. Columns correspond to cells, ordered by Leiden cluster. Dendrogram displays hierarchical clustering. See Table S1 for the full list. (C) UMAP visualization colored by cell type. (D) Inference of chromosomal CNVs on the basis of average expression in windows of 250 analyzed genes. Rows corresponds to cells, ordered by cell type. (E) UMAP visualization colored by CNV score. (F) Dotplot showing the expression of cell markers by each cell type. The size of the dot represents the percentage of cells expressing the gene and the color intensity represents the mean expression in that group. (G) Barplot showing the distribution of cell types obtained in tumors of 7 or 28 days.

To confirm the identity of malignant cells, we inferred copy number variation (CNV) events based on the average expression of 250 genes in each chromosomal region ^4, 28^. We observed large-scale amplifications and deletions in most tumor cells (Figure 2D). We calculated a CNV score for each cell and overlayed it with the UMAP, which revealed eight clusters of cells with high CNV score, including cultured GL261 and GL261-GSCs and cluster 0 (Figure 2E). Figure 2F shows the expression of some of the main marker genes representing each cell type. We also analyzed the spatial topography of gene expression with Visium platform (Table S2). Interestingly, spatial gene expression results confirmed that tumor cell markers shown in Figure 2F, i.e, *Sox2*, *Olig1*, *Nes* and *Pdgfr*, were significantly elevated in the tumor region compared to healthy brain parenchyma. Conversely, astrocyte markers (*GFAP*, *Aqp4* and *Aldh1l1*), neuron markers (*Calb1*, *Slc17a7* and *Gabra1*) and oligodendrocyte markers (*Mbp* and *Mag*) were significantly elevated in the healthy brain parenchyma compared to tumor region (Table S2).

Figure 2G shows the distribution of cell types based on time of tumor progression. Unsurprisingly, most of the cells in 28-day samples had a tumoral origin, whereas we almost did not capture tumor cells in the samples from 7-day tumors. This resulted in a relative detriment in the percentages of all other cell populations at 28 days post-implantation, except for immune cells, whose proportion was increased, indicating high immune infiltration of the tumors. The cell-type identity of each cell cluster along with the number of cells originating from each brain or sample are shown in Table 1 and QC metrics can be found in Suppl. Figure 1.

### Analysis of the transcriptomes of GL261, GL261-GSCs and implanted GL261-GSCs

Given that GBM cells clustered separately depending on their origin (culture in differentiation conditions, culture in stem cell conditions, or implanted into the brain parenchyma), we analyzed the differences between them. Principal component analysis (PCA) revealed that the type of culture and the interaction with the brain microenvironment (brain implantation) are important source of variation (PC1, Figure 3A and Suppl. Figure 3A). We performed Wilcoxon rank sum test to look for potential markers for these populations, with emphasis in the identification of GSC markers in this model, an essential subpopulation for GBM biology and therapy (Figure 3B, Suppl. Figure 2, and Table S3). We found increased expression of cytoskeleton genes, such as *Vim* (vimentin) and *Actb* (beta-actin) in GL261 cells cultured in differentiation conditions. We also found increased expression of several members of the S100 family (including *S100a6*), which present EF-hand domains that bind calcium ions and perform several intracellular and extracellular functions, among which are the regulation of annexin A (AnxA) proteins ^29^, which we also found highly expressed in GL261 cells (including *Anxa2*). AnxA proteins interact with cell membrane components and regulate the structural organization of the cell and intracellular signaling ^30^. Upregulation of some of these proteins may be a reflect of the culture in adherent conditions.

**Figure 3.**
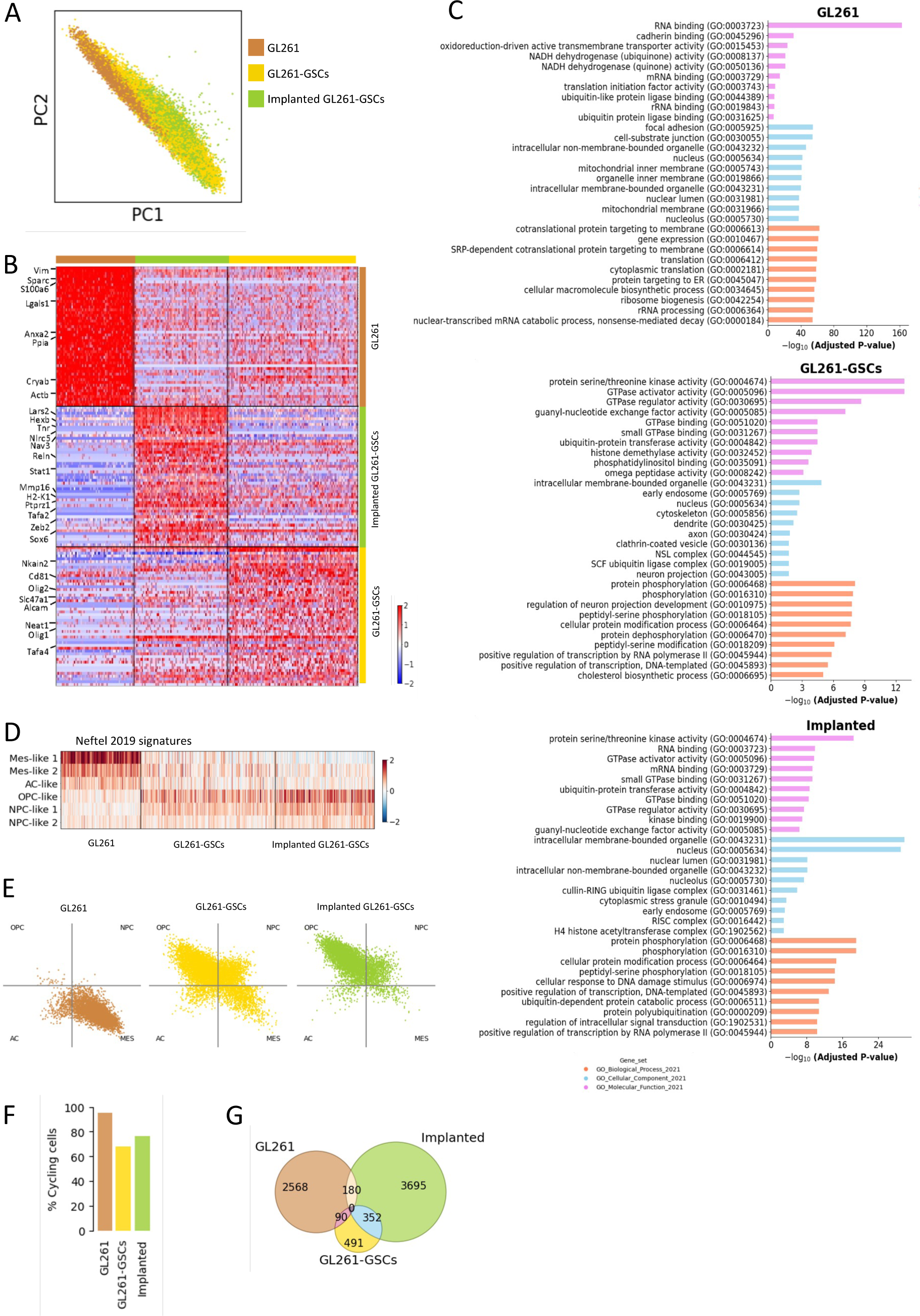
GL261 cells undergo transcriptional changes under different conditions. (A) PCA visualization of malignant cells, colored by cell type. k = 16,757 individual cells. (B) Heatmap showing relative expression of each of the 50 top expressed genes in each culture type. Columns correspond to cultured GL261 and GL261-GSCs and implanted GL261-GSCs. Selected genes are indicated (see Table S4 for the full list). (C) Top 10 enriched terms from gene ontology (GO) gene sets in cultured GL261 and GL261-GSCs and implanted GL261-GSCs compared to the others (see Tables S5, S6, and S7 for full lists). (D) Heatmap showing the meta-module scores described by Neftel et al ^6^. (E) Two-dimensional representation of the cellular states described by Neftel et al.^6^ for each group. Each quadrant corresponds to one cellular state and each dot represents a cell. (F) Barplot showing the abundance of cycling cells, calculated as the sum of cells in S and G2/M phases. (G) Venn diagram showing the overlap between differentially expressed genes across different types. Plotted data are provided in tabular form in Table S7. Differential gene expression was analyzed using Wilcoxon test with a p-value cutoff=0.05

Among overexpressed genes in GL261-GSCs, we found *Nkain2*, *Sema6a, Cdh19* or *Cd81*, all of them proposed as important targets in GBM ^31^. Importantly, the transcription factors *Olig2* and *Olig1*, which showed very high expression in most GL261-GSCs and, to a lesser extent, in implanted GL261-GSCs, were not expressed by GL261 cells (Figure 3B, Suppl. Figure 2), suggesting that they are good markers for GSCs in this model. Also upregulated in GL261-GSCs and, even more, in implanted GSCs was protein tyrosine phosphatase receptor type Z1 (*Ptprz1*) or phosphacan, which has been reported to be expressed by a subset of human GSCs, promoting tumor invasion ^32, 33^.

We used gene set enrichment analysis (GSEA) to identify key pathways and processes (Figure 3C and Tables S4, S5, and S6), and found that most gene ontology (GO) terms upregulated in GL261 cells were related to regulation of the cytoskeleton, cell adhesion, and migration. These results suggest that cell culture in adherent conditions in differentiation medium may promote the expression of a “plastic-attachment signature” where adhesion is an important factor shaping their phenotype. Interestingly, cellular components overrepresented in GL261-GSCs included some neuronal attributes (neuron projection, GO:0043005; dendrite, GO:0030425; axon, GO:0030424), which supports the idea that GL261-GSCs present the potential to differentiate towards a neuron-like lineage. GL261-GSCs upregulated cholesterol biosynthesis, which is linked to cancer development, counting GBM ^34–36^. Cultured and implanted GL261-GSCs were very active in transcription, GTPase activity, and protein modification. Indeed, we scored a set of cell cycle genes from ^37^ and assigned each cell to a cell cycle phase according to the scores obtained. This revealed a similar cell cycle pattern for GL261-GSCs and implanted cells, whereas GL261 samples contained less cells in G1 phase (Figure 3F and Suppl. Figure 3B).

To add additional value to our study, we analyzed GL261 and GL261-GSCs using single-cell methods, a microfluidic-droplet-based method on single cells from 10x (10x cells) and a full-length scRNA-seq using Smart-Seq2 method, based on the separation of cells in microtiter plates by fluorescence activated cell sorting (FACS) ^38^. The number of counts and genes captured per cell were higher using Smart-Seq2 (around 10^4^ genes and 10^6^ total counts), at the cost of sequencing less cells (Suppl. Figure 3C). We detected lower similarity between samples sequenced using Smart-Seq2 compared to that between 10x samples. (Suppl. Figure 3D). 10x nuclei showed the highest dissimilarity with the other technologies used. In the case of GL261-GSCs, the differences between samples sequenced using the same technology were bigger than those observed between GL261 samples, reinforcing the idea that GL261-GSCs are more diverse. We compared differentially expressed genes obtained independently with each technology (Suppl. Figure 4A). From those genes that passed a p-value cutoff of 0.05 in GL261, only 17.51% were shared by the three technologies and 43.69% by two of them, whereas the remaining 38.8% were only detected with one technology. In GL261-GSCs, 14.76% were shared by the three technologies and 26.53% by two of them. 58.71% of the genes were detected only with one technology, with the main contributor to this being Smart-Seq2. We combined information from all three technologies for GO analysis and observed that adding technologies helps identifying different enriched terms within the same gene set, making the analysis more robust, but it does not substantially affect the processes obtained (Suppl. Figure 4B).

Next, we calculated signature scores for the four cellular states that drive the heterogeneity of GBM cells described by Neftel el al. ^6^, i.e., (1) neural-progenitor-like (NPC-like), (2) oligodendrocyte-progenitor-like (OPC-like), (3) astrocyte-like (AC-like), and (4) mesenchymal-like (MES-like). We observed that, whereas GL261 cells span between MES-like and AC-like programs, GL261-GSCs and implanted GL261-GSCs shift towards progenitor states (OPC-like and NPC-like) (Figure 3D). The ‘‘cell-state plot’’ designed by Neftel and colleagues ^6^ summarizes the distribution of cells across these states and their intermediates (Figure 3E). This is in line with recent reports showing a MES-like phenotype of GBM cells grown in two-dimensional culture, with low transcriptional diversity and plenty of cycling cells, and over-representation of developmental states (OPC-, AC-, and NPC-like) in cells cultured in ex vivo human cortical tissue ^33^.

### Effects of the TME on GL261-GSC transcriptional activity

DEG analysis found 2,568 genes unique for GL261 cells, 491 genes unique for GL261-GSCs, and 3,695 genes unique for implanted GL261-GSCs (Figure 3G and Table S7), suggesting that the TME strongly promotes transcriptional heterogeneity, as described in human GBMs ^33^. Implanted GL261-GSCs shared some DEGs with GL261-GSCs (352), indicating that some implanted GSCs retain stem cell properties, and also shared some DEGs with GL261 cultured in differentiation conditions (180), which suggests that GL261-GSCs once implanted in the brain diverge in a great variety of phenotypes, including differentiation programs. Together, these data support the idea that implantation of GL261-GSCs represents a good model to simulate the development of intratumor heterogeneity found in human GBMs. As described above, our results suggest that brain microenvironment importantly affects gene expression in tumoral cells, as judged by the differential gene expression found between implanted and non-implanted GL261-GSCs. Thus, in addition to a downregulation in *Olig2* and *Olig1* expression, we found increased expression of a great number of neuron-related genes, such as *Hexb* (beta-hexosaminidase A subunit beta), *Tnr* (tenascin R), *Nav3* (neuron navigator 3), *Reln* (reelin), and *Tafa2* (TAFA chemokine like family member 2) in implanted GL61-GSCs compared to them in culture (Figure 3B, Suppl. Figure 2, and Table S3). As expected, spatial transcriptomics showed that *Olig2*, *Olig1, Tnr*, *Nav3*, *Reln*, and *Tafa2,* were much more abundant in the tumor area, compared with healthy brain parenchyma (Table S2). Similarly, among the pathways significantly enriched in intracranially implanted GL261-GSCs when compared to cultured GL261-GSCs are many related to synaptic activity and neuronal signaling, such as transmitter-gated ion channel activity (GO:0022824), dendrite membrane (GO:0032590), modulation of chemical synaptic transmission (GO:0050804), inhibitory synapse assembly (GO:1904862), neurotransmitter receptor activity involved in regulation of postsynaptic membrane potential (GO:0099529), neuron projection (GO:0043005), regulation of neurotransmitter receptor activity (GO:0099601), transmitter-gated ion channel activity involved in regulation of postsynaptic membrane potential (GO:1904315), synaptic transmission, GABAergic (GO:0051932) or synapse pruning (GO:0098883) (Table S8), which suggests that the brain TME promotes changes in GL261-GSC transcription towards the development of synaptic activity.

Because of the relevance of the interaction between GBM cells and neurons ^39, 40^, we manually interrogated the expression levels of some of the proposed effectors of this process (Figure 4A). Thus, our study identified the expression of glutamate receptors of the AMPA type (*Gria1*, *Gria2* and *Gria3*), NMDA type (*Grin1*) and Kainate type (*Grik2*) in implanted GL261-GSCs. It should be noted that the expression of AMPA receptors is nearly absent in cultured cells, and it is upregulated in some GL261-GSCs upon transplantation. Tumoral cells also express other proteins associated with neuron-glioma synapsis, such as *Dlg4* (postsynaptic density protein 95, PSD95), *Homer1* (postsynaptic density scaffolding protein PSD-Zip45), and *Nlgn3* (neuroligin-3), one of the key mediators in the pro-tumor effects of neurons on GBM progression ^41^. Interestingly, *Nlgn3* levels were upregulated in tumoral cells by the interaction with the TME, which supports its role as mediator in neuron-glioma synapsis in GL261-GSC GBM model. Some of the GL261-GSCs express the potassium channel KCa3.1 (Figure 4B; *Kcnn4*, also known as IK1 or SK4), with a prominent role in this process ^42^. Implanted GL261-GSCs also express components of tumor microtubes, such as *Gja1* (connexin43), and high levels of *Gap43* (neuronal growth-associated protein 43) (Figure 4B), a key player on the protumor effect of the microenvironment in GBM progression ^43, 44^. While *Gja1* is expressed modestly in all the types of tumor cells, *Gap43* expression importantly increased upon intracranial transplantation (Figure 4B).

**Figure 4.**
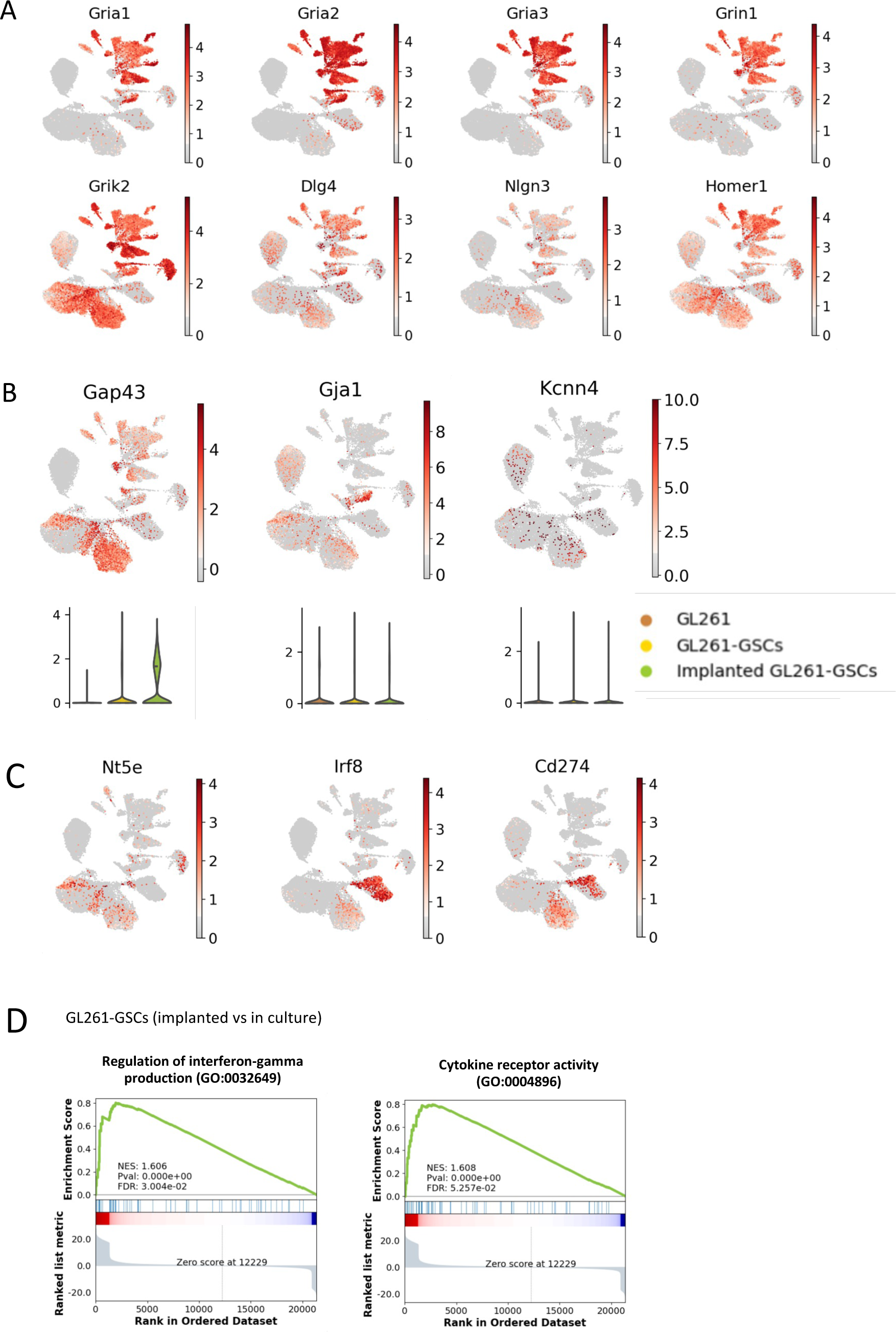
Expression of proteins related to GBM-neuron synapsis and GBM networks communicated through tumor microtubes. (A) UMAPs showing the expression of GBM-neuron synapsis genes. (B) UMAPs and violinplots showing the expression of GBM network genes. (C) Gene set enrichment analysis (GSEA) plot of indicated genesets for genes differentially expressed between implanted and cultured GSCs. NES: Normalized enrichment score; FDR: false discovery rate. (D) UMAPs showing the expression of immune-evasive genes.

Among the genes upregulated in tumor cells by the interaction with the brain microenvironment, we also found increased expression of immune-related genes, including *Nlrc5* (NOD-like receptor family CARD domain containing 5), a transactivator of MHC class I genes, and *H2-K1* (histocompability complex 2 k1) in implanted GL261-GSCs compared to GL261-GSCs (Figure 3B, Suppl. Figure 2, and Table S3). Because GBM cells may acquire an immune evasive phenotype ^45^, we analyzed the expression of the immune evasion regulators *Nt5e* (CD73), *Cd274* (PD-L1), and *Irf8* (interferon regulatory factor 8). We found that *Nt5e* (CD73) was upregulated in implanted and cultured GL261-GSCs as compared to GL261, while the expression of *Cd274* (PD-L1) and *Irf8* was very low in GL261 and GL261-GSCs in culture, but it was upregulated in GL261-GSCs when implanted in the brain for 28 days (Figure 4C). Indeed, spatial transcriptomics showed that the expression of *Nlrc5, Nt5e*, *Cd274*, and *Irf8* was higher in the tumor area, compared with healthy brain parenchyma (Table S2). As a myeloid-specific master transcription factor ^46^, *Irf8* is also expressed by the immune cell cluster (Figure 4C). GSEA analysis showed an increase in “Regulation of interferon-gamma production”, responsible for *Irf8* transcription ^45^, and “Cytokine receptor activity” genesets in GL261-GSCs at 28 days after implantation into the brain when compared to GL261-GSCs in culture (Figure 4D and Table S8).

### The immune microenvironment in the GL261-GSC GBM model

Recomputing Leiden clustering on the 2,427 cells annotated as immune gave 5 clusters (Figure 5A). All 5 clusters expressed the immune cell marker *Ptprc* (CD45, Suppl. Figure 5A). Cluster 3 exhibited higher expression of *Ptprc*, together with lymphocyte markers *Cd4*, *Cd3e*, and *Cd3g* (CD4 T-cell), and Foxp3 (Treg), being thus classified as lymphoid cells, and was almost exclusively composed of cells from the 28-days tumors (Figure 5A-B and Suppl Figure 5A). We did not find expression of B-cell markers. Clusters 0, 1, 2, and 4 were classified as myeloid cells, i.e., tumor-associated microglia/macrophages (TAMs). We observed an increase in the number of cells in G2/M and S phases of the cell cycle in TAMs at 28 days (Figure 5C and Suppl Figure 5B), suggesting an increase in proliferation of myeloid cells throughout GBM development. Scoring gene sets for microglia activation, differentiation, and migration, showed higher activation in immune cells at 28 days, whereas differentiation and migration were reduced (Figure 3D-E). Both stages of tumor development showed similar records of phagocytosis. At 28 days, we observed increased expression of genes associated to major histocompability complex (MHC-II; *Cd74*, *H2-Aa, H2-Ab1, H2-Eb1*), and macrophage markers (*Tgfbi*), concurring with the idea that the myeloid compartment is rich in microglia in the early stages, and in macrophages in the late stages of tumor development (Figure 5F and Suppl. Figure 5C) ^47^ presumably caused by the breakage of the blood-brain barrier observed in human GBMs. Although it is difficult to distinguish tissue resident microglia from blood-derived macrophages, the expression of some genes can help this purpose ^48–50^ (Figure 5G). Microglia has been traditionally identified by expression of *Tmem119*, *P2ry12*, and *Cx3cr1*, whereas blood-derived macrophages express *Tgfbi*, *Mrc1* (mannose receptor C type 1, also called CD206)*, Spp1, Nt5e, S100a4 and Hmox1*, but adapt their transcriptional profile over time (Figure 5H and Suppl. Figure 5D). Expression of markers of GBM-associated macrophages related to immunosuppression, such as *Arg1* (arginase 1), *Mrc1* or *Tgfb1* ^51, 52^ increased throughout tumor development (Figure 5H, Suppl. Figure 5C and 5D and Suppl. Table S9). GSEA revealed upregulation of genes related to TGF-β and Hedgehog signaling, and epithelial-mesenchymal transition at 7 days (Figure 5I and Table S10). TGF-β signaling, triggered by cancer cells, activates pro-inflammatory microglia that have been termed high-grade glioma-associated microglia ^53^. Immune cells at late stage upregulated *Myc* and E2F targets and G2-M checkpoint genes, all indicative of proliferation. IFN-γ, TNF-α signaling, IL-2/STAT5, mTORC1, IL-6/JAK/STAT3, IFN-α, and p53 signaling were also upregulated in 28-days immune cells (Figure 5I and Table S11). mTORC1 activation in microglia leads to the secretion of anti-inflammatory cytokines and immunosuppression ^54^. At both stages, immune cells showed upregulation of genes related to KRAS signaling, inflammatory response, and complement.

**Figure 5.**
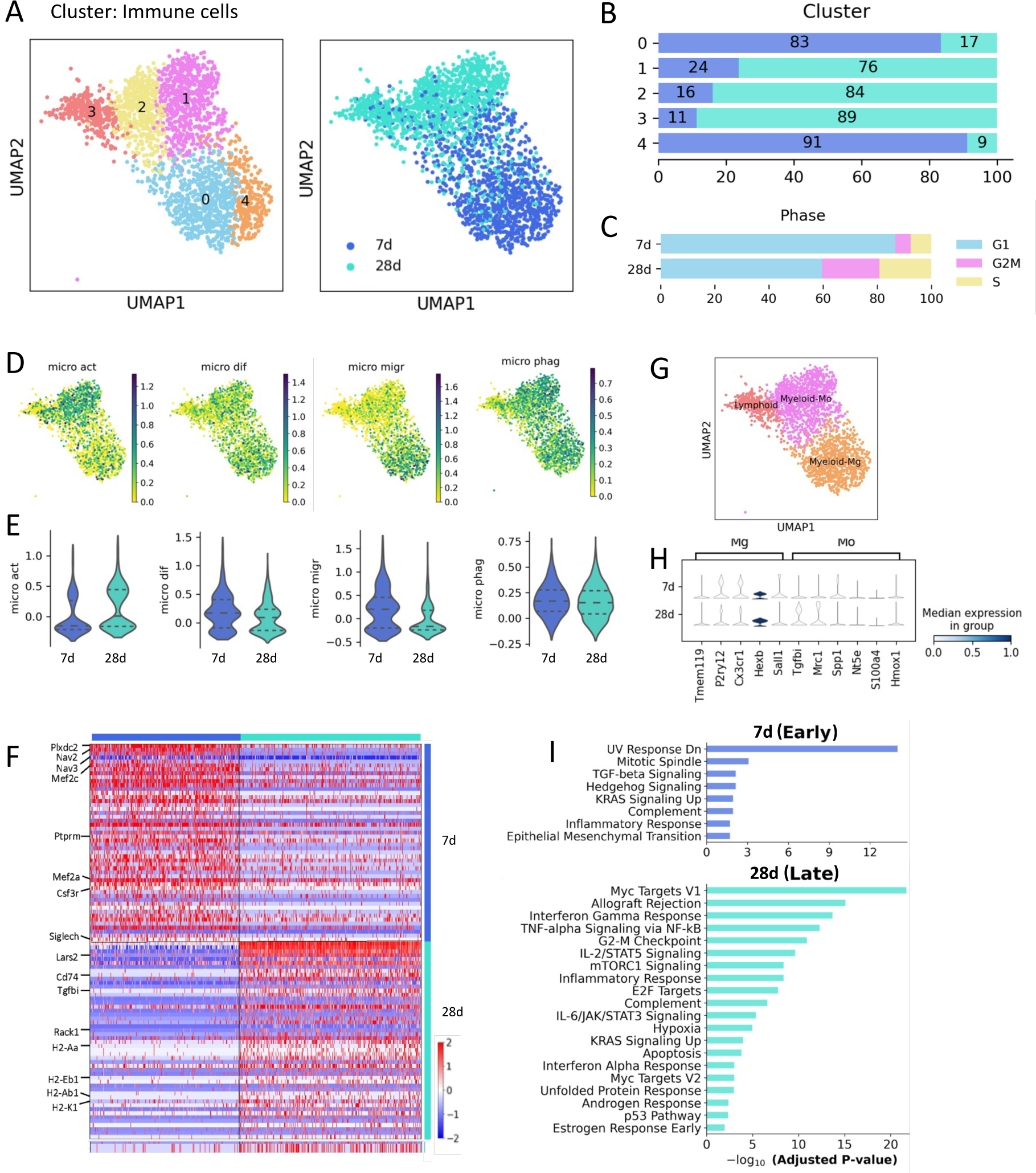
Immune cells in the GL261-GSC model. (A) UMAP visualization of immune cells, colored by Leiden cluster, time of tumor development, and cell type annotations. k = 2,427 individual cells. (B) Barplot showing the proportion of cells at each time point per cluster. (C) Barplot showing the distribution of cells in each cell cycle phase at 7d or 28d. (D) UMAP visualization of microglia activation, microglia differentiation, microglia migration, and microglia phagocytosis scores. (E) Violinplots showing microglia activation, microglia differentiation, microglia migration, and microglia phagocytosis scores at 7 and 28 days. (F) Heatmap showing relative expression of each of the 50 top expressed genes in immune cells at 7 or 28 days. Columns correspond to cells, ordered by time. Selected genes are indicated (see Table S9 for the full list). (G) UMAP visualization of immune cells, colored by cell type annotations. (H) Stacked violinplot showing the expression of microglia and macrophage gene markers in 7d and 28d myeloid cells. (I) Barplot showing all 8 significantly overexpressed terms within the MSigDB Hallmark 2020 geneset in immune cells at 7 days and top 20 overexpressed terms at 28 days (see Tables S10 and S11 for the full lists).

### Cytokine and immune checkpoint landscape in the GL261-GSC GBM model

To provide a tentative map of the factors that mediate the interaction between the developing tumor and the immune system, we analyzed the expression of a panel of cytokines, including chemokines (CCL, CXCL), interleukins (IL), and tumor necrosis factor (TNF) family cytokines and their canonical receptors in all cell clusters (Figure 6A). We found expression of most of them in the immune compartment, remarkably the chemokines *Cxcl10*, *Cxcl9*, *Cxcl16*, and *Ccl25*, *Tgfb1*, *Il1b*, *Tnfsf8*, and *Tnfsf13b*. Malignant cells predominantly expressed the chemokines *Ccl25* and *Cxcl10*, Colony stimulating factor 1 (CSF-1; *Csf1)*, *Il7*, *Il33*, *Il34*, *Il17d*, *Tnfsf10*, and *Tnfsf13b*. *Ccl25* encodes an important cytokine in tumor progression ^55^ highly expressed in all clusters in our data, and especially in implanted GL261-GSCs. However, its canonical receptor, CCR9, was scarcely expressed only by endothelial cells. *Cxcl10* was mainly expressed by tumor cells, and to a lesser extent by other TME cells, such as immune cells, astrocytes, endothelial cells and OPCs. CXCL10 is secreted in response to IFN-γ, whose production is upregulated in implanted GL261-GLSCs (Figure 4D). Intriguingly, the expression of *Cxcr3*, the canonical receptor of CXCL10, was not found in our data. *Csf1* was expressed by tumor cells and other TME cells, such as astrocytes and endothelial cells, while its receptor, *Csf1r*, was highly expressed by immune cells. IL-34 is another ligand for CSF-1R, which regulates microglia and macrophages, and it has the ability to interact with PTPRZ ^56^, whose transcript we found within the most overexpressed in GL261-GSCs, specially in implanted GL261-GSCs (Figure 3B and Suppl. Figure 2). GL261-GSCs also expressed *Il17d*, which can suppress the activity of CD8^+^ T cells by regulating dendritic cells ^57^ and *Tnfsf10* and *Tnfsf13b,* which encode two important proteins in cancer biology, TNF-related apoptosis-inducing ligand (TRAIL; ^58^ and TNF- and APOL-related leukocyte expressed ligand 1 (TALL-1), also called B-cell activating factor (BAFF; ^59^). Among the receptors for these proteins, in this study we found a modest expression of *Tnfrsf10b* and *Tnfrsf13b* and *Tnfrsf13c* in tumor cells and astrocytes, and *Tnfrsf13b* and *Tnfrsf13c* in immune cells. Endothelial cells showed high expression of *Cxcl12*, as previously described ^47^, *Il34*, and *Tnfsf10*. Oligodendrocytes showed strikingly high expression of *Il33*, which is known to be released upon CNS injury as an alarmin that acts on local astrocytes and microglia to induce chemokines for monocyte recruitment ^60^, while being implicated in oligodendrocyte maturation ^61^. Transforming growth factor beta 1 (*Tgfb1*) transcript was found in all cell types, accompanied by high expression of its receptors *Tgfbr1* and *Tgfbr2* in endothelial, tumor and immune cells, and *Tgfbr3* in endothelial cells. Strangely, we did not find expression of the interferon gamma (*Ifng*) transcript, although expression of its receptors *Ifngr1* and *Ifngr2* was found in all cell types, especially immune cells.

**Figure 6.**
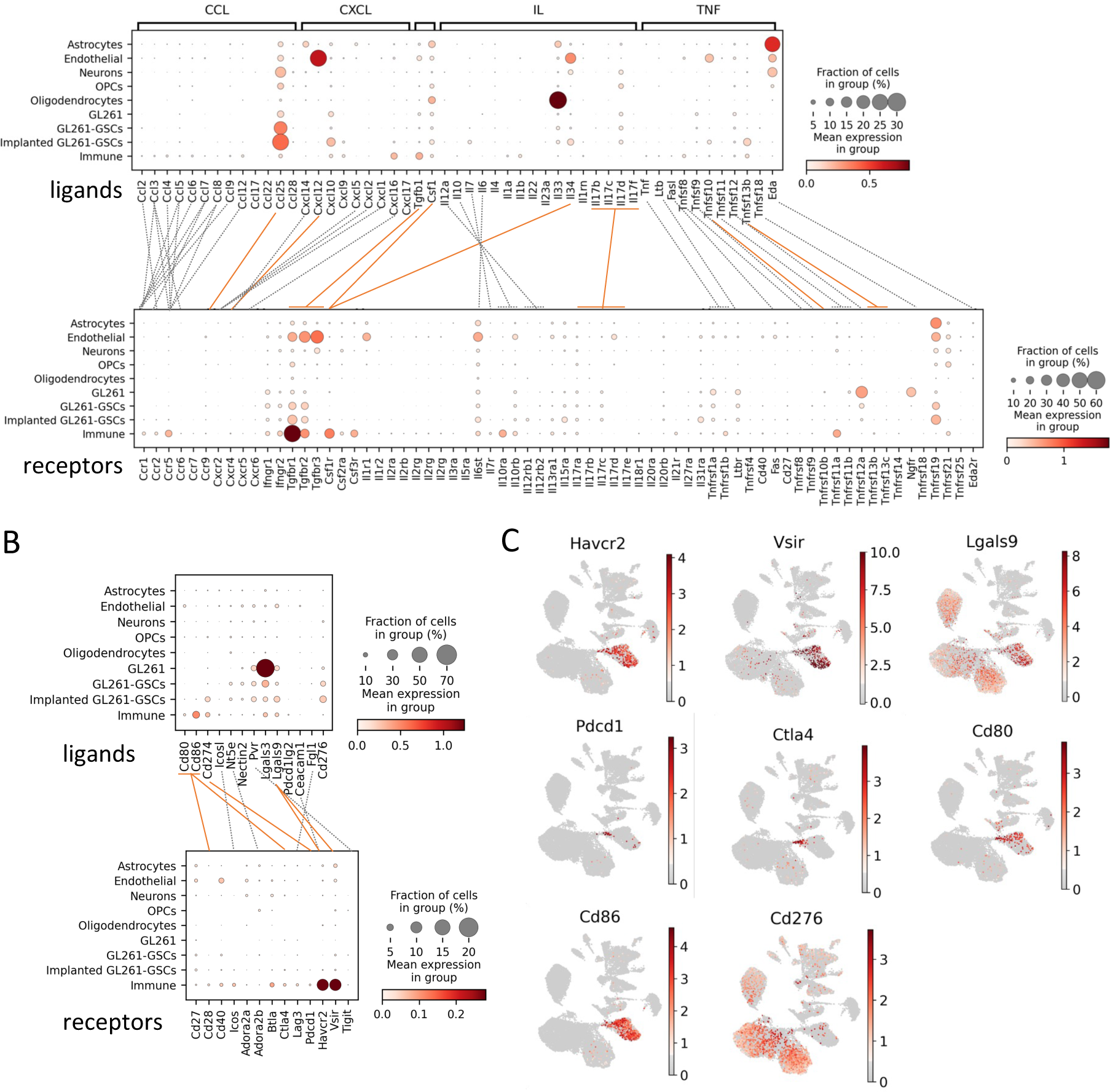
Expression of cytokines and checkpoint molecules. (A) Dotplot showing expression of cytokines and receptors per cluster. Orange lines indicate interactions described in the text; gray dotted lines indicate other described interactions. (B) Dotplot showing expression of checkpoint ligands and receptors per cluster. Orange lines indicate interactions described in the text; gray dotted lines indicate other described interactions. (C) UMAPs showing the expression of selected genes.

Because of the relevance of checkpoints inhibitor in immunotherapy, we analyzed the expression of the main checkpoint ligands and receptors in all the cell clusters (Figure 6B). Immune cells showed expression of checkpoint receptors, with strikingly high levels of TIM-3 (*Havcr2*) and VISTA (*Vsir*) in TAMs (Figure 6B and C). Checkpoint ligands were expressed by tumor cells and immune cells, accordingly with recent studies that show the expression of checkpoint molecules by myeloid cells ^47, 62^. TIM-3 and VISTA ligand, Galectin-9 (*Lgals9*), was expressed in both tumor and immune cells. We observed low levels of PD-1 transcripts (*Pdcd1*), restricted, as expected, mostly to the T cell cluster. PD-L1 transcript (*Cd274*) levels were more prominent and were mostly present in myeloid immune cells and in tumor cells. PD-L2 (*Pdcd1lg2*) levels were very low, maybe due to the low abundance of dendritic cells in our data. Expression of the co-stimulatory molecule *Cd276* (B7-H3) increased in GSCs, especially after transplantation. *Ctla4* expression is low and restricted to T cells, whereas its B7 ligands (*Cd80*, and mainly *Cd86*) were also found in myeloid cells.

Of note, spatial transcriptomic results showed that expression of most of the immune-related genes found in this study was significantly higher in the tumor area compared to healthy brain parenchyma (Table S2). This is the case for markers of different immune cell types (*Ptprc, Cd4*, *Cd3e*, and *Cd3g, Cd74*, *H2-Aa, H2-Ab1, H2-Eb1, Tmem119, Cx3cr1, Mrc1, Nt5e, S100a4, Hmox1, Arg1*, *Mrc1* or *Tgfb1*), cytokines and receptors (*Ccl25, Cxcl10,* IFN-γ, *Csf1, Csf1r*, *Il17d*, *Tnfsf10*, *Tnfsf13b, Tnfrsf10b, Tnfrsf13b*, *Tnfrsf13c, Tgfb1*, *Tgfbr1, Ifng*, *Ifngr1* and *Ifngr*) and checkpoints and receptors (*Havcr2, Vsir, Lgals9, Cd274, Pdcd1lg2, Cd276, Ctla4 and Cd86)*.

### snRNA-Seq analysis of TMZ and Tat-Cx43_266-283_ treatments in GL261-GSC GBM model

Following the same snRNA-Seq protocol, we analyzed the brain tumors and their microenvironment in mice treated with temozolomide (TMZ), the standard of care for GBM patients, or with the cell penetrating peptide Tat-Cx43_266-283_ (Tat-Cx43) ^21, 22, 24^ at 7 and 28 days post-implantation, as described in Figure 1A. Clusters were classified by treatments, annotations (Figure 7A), time and samples (Suppl. Figure 6A) as previously described. Although extensive information can be obtained from these results, we focused on those results highlighted as more prominent for this preclinical model in previous sections.

**Figure 7.**
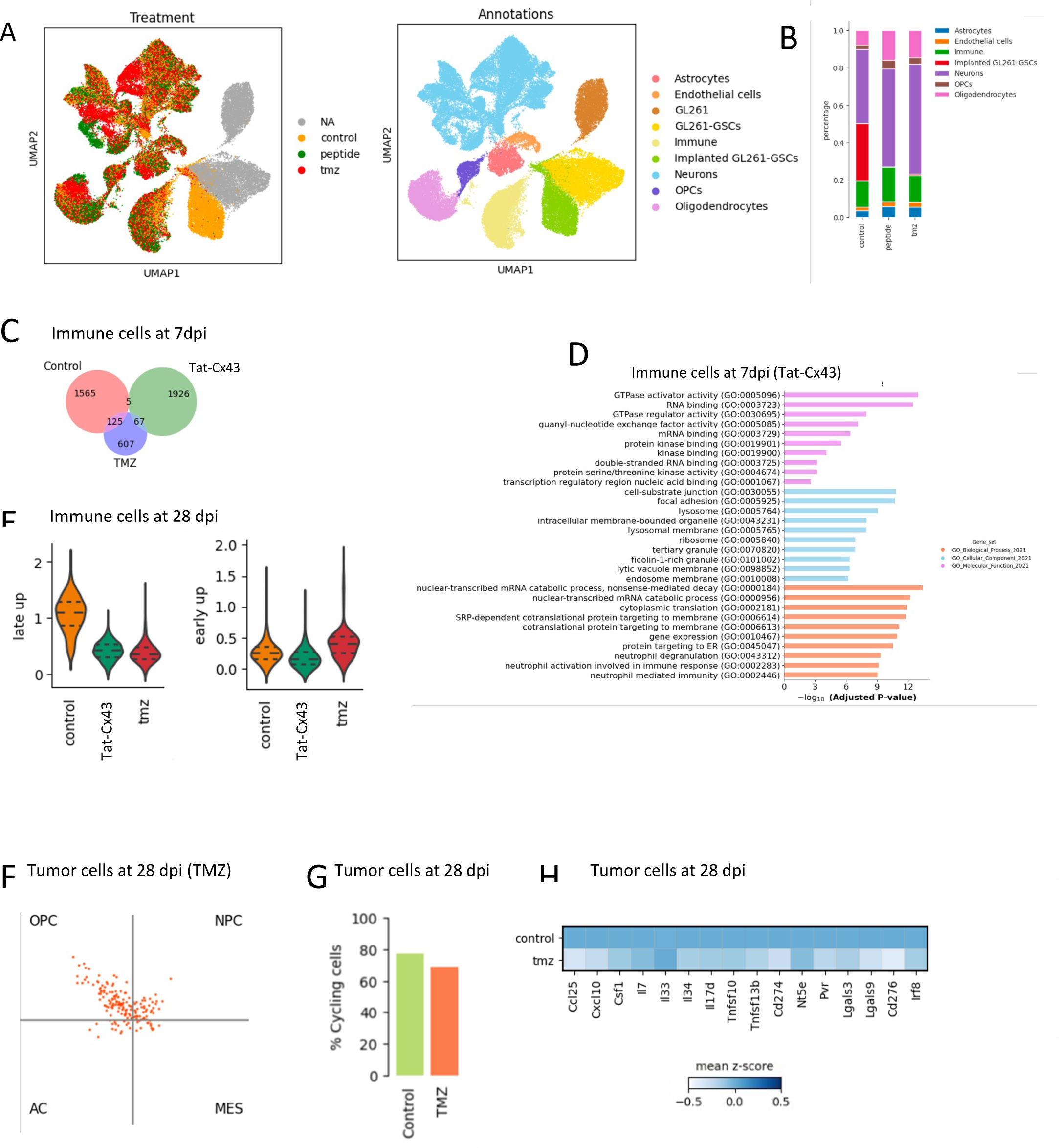
Effect of Tat-Cx43 and temozolomide (TMZ) treatment on immune and tumor cells at 7- and 28-days post-implantation (dpi). (A) UMAP visualization of all cells collected, colored by treatment and annotations. (B) Barplot showing the distribution of cell types obtained in untreated and treated animals. (C) Venn diagram showing the overlap between differentially expressed genes across treatments in the immune cell cluster at 7 dpi. Differential gene expression was analyzed using Wilcoxon test with a p-value cutoff=0.05. See Table S12 for full list. (D) Top 10 enriched terms from gene ontology (GO) gene sets in immune cell cluster from Tat-Cx43-treated mice compared to the control (see Table S13 for full list). (E) Violinplots showing early (high levels in controls at 7 days) and late (high levels in controls at 28 days) immune scores in immune cells at 28 days post-implantation under different treatments. (F) Two-dimensional representation of the cellular states described by Neftel et al. ^6^ in tumor cells of temozolomide-treated mice at 28 dpi. Each quadrant corresponds to one cellular state and each dot represents a cell. (G) Barplot showing the abundance of cycling cells, calculated as the sum of cells in S and G2/M phases, in control and TMZ-treated tumor cells at 28 dpi. (H) Heatmap showing the expression of selected cytokines and checkpoint molecules in control and TMZ-treated tumor cells at 28 dpi.

Given the low numbers of malignant cells obtained from 7-day samples (Suppl. Figure 6A), we explored the immune cell cluster at this stage. As shown in Figure 7C, the treatment with Tat-Cx43_266-283_ peptide extensively affected the transcription in immune cells at 7 days post tumor implantation (see Table S12 for the full list of DEGs). Indeed, 1,926 genes were differentially expressed by immune cells in Tat-Cx43_266-283_-treated animals, while 607 were modified in TMZ-treated animals. Only 67 genes were commonly affected by TMZ and Tat-Cx43_266-283_. The analysis of top 10 enrichment biological processes, cellular components, and molecular functions from gene ontology (GO) unveiled interesting GTPase, kinase, translation, transcriptional, and immune-related activity of immune cells in brain tumors treated with Tat-Cx43_266-283_ (Figure 7D and Table S13). These analyses showed only molecular functions, mostly related to immune system, significantly modified by TMZ (Suppl. Figure 6B and Table S14). By comparing the expression of genes in immune cells in untreated tumor-bearing mice, at 7 vs 28 days post-implantation, we obtained an early (7 days) and late (28 days) immune score. Interestingly, the treatment with Tat-Cx43_266-283_ and TMZ for 28 days, decreased the expression of those genes elevated in untreated animals at 28 days (late up) and showed an expression pattern more similar to that found at 7 days (early up) in untreated animals (Figure 7E). Suppl. Figure 6C shows the reduction in the expression of some checkpoint molecules and immune modulators, such as *Hexb, Lars2, Rack1, H2-K1, Mrc1, Tgfb1, Cd74, Lglas9 or Nav3* promoted by Tat-Cx43_266-283_ and TMZ in immune cell cluster at 28 days post tumor implantation.

Because of the low numbers of malignant cells obtained from peptide-treated 28-day samples, we could only study differences in malignant cells from control and TMZ-treated 28-day samples. Figure 7F shows that, upon treatment with TMZ for 28 days, tumor cells mostly fall into OPC and, to a lesser extent, NPC categories in the cellular-state map from Neftel et al. ^6^. As expected, the analysis of cell cycle by snRNA-Seq showed that tumor cell proliferation was decreased by TMZ treatment (Figure 7G). Interestingly, the expression of key genes involved in immune evasion, such as *Irf8*, *Cd274*, *Lgals9*, *Havcr2*, and *Cd276* were downregulated by TMZ in tumoral cells (Figure 7H and Table S15). Furthermore, TMZ significantly decreased the expression of key genes related to neuron-glioma synapsis, such as *Gap43*, *Gja1*, *Gria3*, *Grik2* and *Homer1* (Table S14) in tumoral cells. However, some genes related to bad prognosis were upregulated in tumoral cells treated with TMZ, including as *Ahnak*, *Atrx*, *Cd44*, *Jarid2*, *Plcd4*, *Cdk8* and *Myc* (Suppl. Figure 6D).

## Discussion

Preclinical models are essential to advance our understanding of GBM biology and to improve their treatment. However, a thorough characterization is required to identify the cellular and molecular targets that can be studied and, regarding the application of these results, the type of human GBM which will benefit from these results. In this study, we provide a comprehensive characterization of the preclinical GL261-GSC GBM model, in which tumors are originated by murine GSCs intracranially implanted in immunocompetent mice. We chose this model because it emulates the origin of human brain tumors from GSCs ^63, 64^ and allows the study of the GBM immune system and its evolution throughout tumor development. Our snRNA-Seq data showed that GL261-GSCs recapitulate some key human GSC attributes, cellular pathways, and processes. More importantly, once implanted into the brain, GL261-GSCs diverge and give rise to heterogenous GBM cells, which share the expression of some genes with differentiated GL261 cells and GL261-GSCs, but mainly express unique genes, indicating that these tumors are heterogenous and include stem-like and differentiated GBM cell subpopulations.

Interestingly, we found a bidirectional interdependence between tumoral cells and brain microenvironment, which is very similar to that described in human GBM ^33^. Although these results can be mined with a great variety of purposes, we focused on the transcriptional changes promoted by the brain microenvironment on tumoral cells and those promoted by tumoral cells on the immune system. These analyses unveiled the expression of key GBM targets that can be studied in the GL261-GSC GBM model. Furthermore, we found similarities with a subtype of human GBM, paving the way to translate the preclinical results generated from the GL261-GSC GBM model into clinical trials in this specific human GBM subtype.

### Feasibility of the GL261-GSC GBM model to study GBM-neuron synapsis and GBM networks

Increasing evidence has demonstrated that GBM progression is robustly regulated by neuronal activity, which stimulates synaptic activity in GBM cells ^39, 40, 42, 65^. Our results show that among the pathways significantly enriched in intracranially implanted GL261-GSCs when compared to cultured GL261-GSCs are many related to synaptic activity and neuronal signaling, which suggests that the brain TME promotes changes in transcription towards the development of synaptic activity. Indeed, implanted GL261-GSCs expressed diverse ionotropic glutamate receptors, which might activate intercellular calcium signaling networks to orchestrate GBM cell growth, invasion, and drug resistance as it has been previously shown_39,40,42,65._

Neuronal input facilitates the formation of tumor microtubes, which are ultra-long membrane tube protrusions that connect single tumor cells to create functional and communicating multicellular networks by intercellular Ca^2+^ waves. The neuronal growth-associated protein 43 (*Gap43*) is important for microtube formation and function, and drives microtube-dependent tumor cell invasion, proliferation, interconnection, and radioresistance ^43, 44^. Microtube-associated gap junctions formed by connexin43 (*Gja1*) also contributes to communication in the multicellular network. The increased expression of *Gap43* found in GL261-GSCs upon intracranial transplantation, together with *Gja1* (Cx43) expression, is compatible with a network of communicating tumor microtubes in the GL261-GSC GBM model. In this regard, it has been recently shown that the potassium channel KCa3.1 (*Kcnn4*) is present in a small population of human GBM cells and is responsible for rhythmic Ca^2+^ oscillations within the connected network that sustain tumor growth ^42^. Our study showed that some of the implanted GL261-GSCs expressed channel KCa3.1 (*Kcnn4*). Furthermore, the high levels of *Tgfb1* mRNA found in immune cells in the GL261-GSC GBM model might contribute to the formation of microtubes ^66^. Implanted GL261-GSCs were predominately enriched for the oligodendrocyte precursor-like (OPC-like)/neural precursor-like (NPC-like) cell states, which coincides with the most migratory GBM subpopulation described by Venkataramani et al. ^65^. Therefore, the GL261-GSC GBM model might be useful to study the mechanism by which GBM-neuron synapsis are developed and to carry out preclinical studies of drug candidates designed to target key candidates, such as GAP43, ionotropic glutamate receptors, neuroligin-3 ^41^ or KCa3.1, specifically in GBM cells.

### Intracranially-injected GL261-GSCs develop an immune evasive phenotype

Previous studies showed that GBM cells may acquire an immune evasive phenotype via epigenetic immunoediting ^45^, a process in which immune attack promotes transcriptional changes in tumoral cells that are stabilized and selected in those cells with increased immune evasive qualities, resulting in highly immune evasive and transcriptionally altered descendants. We found that key genes whose methylation is erased and are immune evasion regulators, such as *Nt5e* (CD73), *Cd274* (PD-L1), and *Irf8* are expressed by GL261-GSCs upon brain transplantation. In fact, most of them are barely expressed in GL261-GSCs but are upregulated when implanted in the brain for 28 days, indicating that they are expressed in response to TME. CD73 is a critical component in the formation of an immunosuppressive microenvironment in cancer and its expression correlates with bad prognosis in distinct types of tumors ^67^. CD73 is secreted in GBM cell exosomes ^68^ and has been identified as a relevant target to improve immune checkpoint therapy in GBM ^69^. *Cd274*, also known as programmed death-ligand 1 (PD-L1) is one of the best immune scape mediators and the development of anti-PD1/PD-L1 antibodies is a hot topic in cancer immunotherapy ^67^. Upon binding to PD1, it activates downstream signaling pathways and inhibits T cell activation. Irf8 is the key master transcription factor for immune evasion, and it is upregulated in tumor cells following immune attack via IFN- γ -mediated activation ^45^. Our study unveiled “Regulation of interferon-gamma production” among the gene sets upregulated in GL261-GSCs after implantation into the brain when compared to GL261-GSCs in culture. Altogether, these results indicate that the in vivo immune attack triggers significant transcriptional changes in GL261-GSCs to promote an immune evasive phenotype.

In line with the immune evasion developed by tumor cells in response to TME, the expression of cytokine and cytokine receptors found in this study suggest that the development of these tumors reshapes the TME, with a shift towards a supportive immune system where TAMs are the main allies. Thus, *Csf1r, Arg1, Mrc1 or Tgfb1*^51, 52^ are among the markers of immunosuppressive and tumor supportive myeloid cells highly expressed by the immune cluster, mainly at late stages of tumor development. Therefore, this is an interesting preclinical model to study the development of immune evasion in GBM cells promoted by the brain microenvironment and to explore therapies against crucial targets for this process in tumor cells, such as *Nt5e* (CD73)*, PD-L1*(Cd274) or *Irf8,* and in immune cells, such as *Csfr1, Arg1, Mrc1 or Tgfb1*.

### The GL261-GSC model resembles a specific subtype of human GBM with a heterogenous immune population

One of the hallmarks of human GBM is the infiltration of immune cells that may constitute up to a 30% of the cells that integrate these tumors. Spatial transcriptomics showed that the expression of most of the immune-related genes was significantly higher in the tumor area compared to healthy brain parenchyma, indicating that immune cells infiltrated brain tumors in the GL261-GSC GBM model, as described in human GBMs. Three novel human GBM subtypes with significantly different TME compositions have been recently proposed based on the analysis of more than 800 human GBM samples ^3^. These subgroups were defined as TMELow (Immune-Low), TMEMed (heterogenous immune populations) and TMEHigh (Immune-High), comprising 24%, 46% and 30% of human GBMs, respectively. Only those patients classified as TMEHigh showed a significant improved survival after immunotherapy, which highlights the relevance of patient stratification to rationally select patients who might benefit most from treatment. By comparing these human GBM subtypes with the GL261-GSC GBM murine model, we found substantial similarities with TMEmed subtype, the most frequent human GBMs ^3^. TMEmed GBMs are characterized by heterogeneous immune populations, enrichment in endothelial cell gene expression profiles and in pathways related to neuronal signaling. These and other TMEmed GBM features, such as the downregulation of the B lymphocyte chemoattractant and TLS marker, *Cxcl13*, are found in the GL261-GSC GBM murine model. More relevant for immunotherapy, *Pdcd1* (PD1), *Ctla4* and *Lag3* are scarcely expressed in TMEmed GBM subtype, as well as in the GL261-GSC GBM model, which is in contrast with the high level of expression of these immune checkpoints in TMEhigh GBM subtype tumors. Retrospective studies showed that only TMEhigh GBMs have a positive response to anti-PD1 immunotherapy ^3^, which agrees with the high levels of PD1 expression. Interestingly, GBM models derived from differentiated GL261 cells are highly responsive to anti-PD1 and anti-CTLA4 immunotherapy ^70^, indicating high levels of PD1 and CTLA4 in differentiated GL261 model. These data support the immune evasive behavior of the GL261-GSC subpopulation and its descendants described above and suggest that immune evasion might be a stem cell specific feature. Importantly, other key targets for immunotherapy, such as PD-L1, TIM-3 or B7-H3 are highly expressed in the GL261-GSC GBM model, which coincides with TMEmed GBM subtype ^3^, indicating the utility of GL261-GSC GBM model as a preclinical model to test therapies against these immunotherapy targets and support the translation of GL261-GSC GBM results in TMEmed GBM subtype.

### Study limitations and future perspectives

Among the limitations of this study is that it shows mRNA levels, therefore the data should be confirmed at the protein level. As described in this study, scRNA-seq and other technologies may provide complementary information to snRNA-Seq, as each technology provides some non-overlapping additional information. Finally, although most data analyzed through the manuscript is robust, the low number of nuclei in some clusters under specific conditions, for example GBM cells after 7 days post-implantation, is limited and these results should be interpreted with caution.

Overall, this study will serve to explore the utility of the GL261-GSC GBM model to study specific GBM targets and potential treatments with a strong rationale and better options for successful clinical translation. In conclusion, this work provides crucial information for future preclinical studies in GBM improving their clinical application.

## Materials and methods

### Animals

C57BL/6 mice were shipped from Charles River to the animal facility of the University of Salamanca at INCYL (SEA-INCYL). Animals were housed on a 12h/12h light/dark cycle and provided with food and water ad libitum. Mice were maintained singly from the start of the experiments, and were monitored for signs of humane endpoints daily, including changes in behavior and weight. All animal procedures were approved by the ethics committee of the University of Salamanca and the Junta de Castilla y León (Spain) and were carried out in accordance with European Community Council directives (2010/63/UE), and Spanish law (RD 53/2013 BOE 34/11370–420, 2013) for the use and care of laboratory animals.

### Cells

GL261 cells were grown adherently in differentiation medium containing DMEM supplemented with 10% fetal calf serum (FCS), dissociated using trypsin/EDTA, and split to convenience. Neurospheres (GL261-GSCs) were obtained from GL261 adherent cultures as previously described ^20^ and cultured in stem cell medium containing Dulbecco’s modified Eagle’s medium (DMEM)/Nutrient Mixture F-12 Ham supplemented with 1% Minimum Essential Medium-Non-Essential Amino Acids (MEM-NEAA), 3.9 mM glucose, 1 mM L-glutamine, 0.07% β-mercaptoethanol, 121.8 μg/mL bovine serum albumin (BSA), 1% B-27 supplement, 0.5% N-2 supplement, 10 ng/ml epidermal growth factor (EGF), and 10 ng/ml basic fibroblast growth factor (b-FGF). Neurospheres were dissociated using Accutase and subcultured at a density of 10^4^ cells/ml every 8-10 days. GL261-GSCs were stably transfected with pcDNA3.1-mCherry plasmid (a kind gift from C. Naus) using Lipofectamine 2000 Transfection Reagent according to manufacturer’s instructions. Cells were plated at low density and mCherry^+^ colonies were selected and amplified. All culture media were supplemented with 50 U/ml penicillin G, 37.5 U/ml Streptomycin and 0.23 μg/ml Amphotericin B to avoid bacterial and fungal contamination. Cells were maintained at 37°C in an atmosphere of 95% air/5% CO^2^ and with 90-95% humidity.

### Intracranial implantation of glioma cells

mCherry-GL261-GSCs were injected into 8-week-old C57BL/6 mice as previously described ^21^. An equal number of males and females was used. Mice were anesthetized by isoflurane inhalation, placed on a stereotaxic frame, and window-trephined in the parietal bone. A unilateral intracerebral injection to the right cortex was performed with a Hamilton microsyringe. 1 μl of physiological saline containing 5,000 cells was injected at the following coordinates: 1 mm rostral to lambda, 1 mm lateral, and 2 mm deep. To minimize the inflammatory response from damaged brain tissue due to the needle injection, tumoral cells were slowly injected into the brain and the needle was held in place for an additional 2 minutes before removal. Cellular suspensions were kept on ice while the surgery was being performed and allowed to temperate for five minutes once loaded into the microsyringe. At the indicated times, mice were anesthetized with pentobarbital (120 mg/kg, 0.2 ml) and transcardially perfused with 15 ml of physiological saline. Brains were removed, fresh-frozen in liquid nitrogen, and kept at −80 °C until used.

### Treatments

Synthetic peptides (>95% pure) were obtained from GenScript. YGRKKRRQRRR was used as the Tat sequence, which enables the cell penetration of peptides. The Tat-Cx43_266-283_ sequence was Tat-AYFNGCSSPTAPLSPMSP (patent ID: WO2014191608A1). 100 μM Tat-Cx43_266-283_ or 250 μM TMZ were intracranially injected in 1 μl of physiological saline together with the cells. For 28-day experiments, 4 nmol/g Tat-Cx43_266–283_ or 10 nmol/g TMZ were intraperitonially injected twice per week, starting on day 8 and until the end of the experiment. An equivalent amount of saline was injected to control mice.

### Tissue dissociation and sample preparation

A tissue piece was collected from each brain. Each sample included tumor core and peritumoral space. Nuclei suspensions from frozen tissue were obtained using a density gradient medium as described (https://www.protocols.io/view/nuclei-isolation-prep-and-protocol-5qpvoneedl4o/v1). Tissue pieces were mechanically digested in ice-cold buffer containing protease and RNAse inhibitors using a Dounce homogenizer and incubated in 0.35% IGEPAL® CA-630. Homogenized solutions were filtered using a 40 μm-pore cell strainer and carefully added to ultracentrifuge tubes with layered 40% and 30% Iodixanol solutions. Tubes were centrifuged at 10,000 *g* for 18 minutes and nuclei were collected from the interface between 40% and 30% Iodixanol solutions. Nuclei suspensions from fresh cells were obtained following 10xGenomics protocol (https://www.10xgenomics.com/products/nuclei-isolation). Cells and neurospheres were dissociated using Accutase and lysed in lysis buffer containing 0.3% IGEPAL® CA-630. Nuclei were counted using a Countess Automated Cell Counter and nuclei quality was assessed in an inverted microscope. Cells and nuclei were loaded with a target output of 6,000 nuclei per sample.

### FACS

All protocols used in this study are described in detail elsewhere ^71^, including preparation of lysis plates, FACS sorting, cDNA synthesis using the Smart-Seq2 protocol ^72, 73^, library preparation using an in-house version of Tn5 ^74, 75^, library pooling and quality control, and sequencing. For further details please refer to https://doi.org/10.17504/protocols.io.2uwgexe. Single-cell suspensions were sorted by a SONY SH800 Cell Sorter. All events were gated with forward scatter-area (FCS-A)/side scatter-area (SSC-A) and FCS-height (FCS-H)/FCS-width (FCS-W). Single cells were sorted in 96-well plates containing 4 μL lysis buffer (4U Recombinant RNase Inhibitor, 0.05% Triton X-100, 2.5 mM dNTP mix, 2.5 μM Oligo-dT30VN (5ʹ-AAGCAGTGGTATCAACGCAGAGTACT30VN-3ʹ), spun down for 2 min at 1,000 × *g*, and snap frozen. Plates containing sorted cells were stored at −80°C until processing. Reverse transcription and PCR amplification were performed according to the Smart-seq2 protocol described previously ^72^. In brief, 96-well plates containing single-cell lysates were thawed on ice followed by incubation at 72°C for 3 min and placed immediately on ice. Reverse transcription was carried out after adding 6 μL of reverse transcription-mix (100 U SMARTScribe Reverse Transcriptase, 10 U Recombinant RNase Inhibitor, 1× First-Strand Buffer, 8.5 mM DTT, 0.4 mM betaine, 10 mM MgCl_2_, and 1.6 μM TSO (5ʹ-AAGCAGTGGTATCAACGCAGAGTACATrGrG+G-3ʹ) for 90 minutes at 42°C, followed by 5 minutes at 70°C. Reverse transcription was followed by PCR amplification. PCR was performed with 15 μL PCR mix (1× KAPA HiFi HotStart ReadyMix), 0.16 μM ISPCR oligo (5ʹ-AAGCAGTGGTATCAACGCAGAGT-3ʹ), and 0.56 U Lambda exonuclease according to the following thermal-cycling protocol: (1) 37°C for 30 min; (2) 95°C for 3 min; (3) 21 cycles of 98°C for 20 s, 67°C for 15 s, and 72°C for 4 min; and (4) 72°C for 5 min. PCR was followed by bead purification using 0.7× AMPure beads, capillary electrophoresis, and smear analysis using a Fragment Analyzer. Calculated smear concentrations within the size range of 500 and 5,000 bp for each single cell were used to dilute samples for Nextera library preparation as described previously ^73^.

### Microfluidic droplet single-cell analysis

Single cells or nuclei were captured in droplet emulsions using the Chromium instrument (10x Genomics) and scRNA-seq libraries were constructed as per the 10x Genomics protocol using GemCode Single-Cell 5’ Bead and Library kit. Single nuclei were processed using the 10x Multiome ATAC + Gene Expression kit. All reactions were performed in the Biorad C1000 Touch Thermal cycler with 96-Deep Well Reaction Module. The number of cycles used for cDNA amplification and sample index PCR was determined following 10x guidelines. Amplified cDNA and final libraries were evaluated on a TapeStation System using a High Sensitivity D5000 ScreenTape. The average fragment length of the libraries was quantitated on a TapeStation System, and by qPCR with the Kapa Library Quantification kit for Illumina. Each library was diluted to 2 nM and equal volumes of up to 8 libraries were pooled for each sequencing run. Pools were sequenced with the number of cycles indicated by 10x guidelines. A PhiX control library was spiked in at 1%. Libraries were sequenced on a NovaSeq 6000 or a NextSeq 2000 Sequencing System (Illumina).

### Visium spatial transcriptomics

Visium spatial transcriptomics were carried out as previously described by Janesick et al. (2022; doi: https://doi.org/10.1101/2022.10.06.510405). Briefly, after intracranial tumor implantation, frozen brain sections were obtained with a cryostat and processed for methanol fixation, hematoxylin eosin staining and imaging of tissue for use with 10x Genomics Visium Spatial protocols, following manufacturer’s instructions.

### Sequencing data extraction and pre-processing

Sequences were de-multiplexed using bcl2fastq v2.19.0.316. Reads were aligned to the mm10plus genome using STAR v2.5.2b with parameters TK. Gene counts were produced using HTSEQ v0,6,1p1 with default parameters, except ‘stranded’ was set to ‘false’, and ‘mode’ was set to ‘intersection-nonempty’. Sequences from the microfluidic droplet platform were de-multiplexed and aligned using CellRanger v2.0.1, with default parameters. Gene count tables were combined with the metadata variables using the Scanpy Python package v.1.4.2. We removed genes that were not expressed in at least 3 cells and then cells that did not have at least 250 detected genes. For FACS we removed cells with fewer than 5,000 counts, and for the droplet method we removed cells with fewer than 2,500 unique molecular identifiers (UMIs). The data was then normalized using size factor normalization such that every cell has 10,000 counts and log transformed. We computed highly variable genes using default parameters and then scaled the data to a maximum value of 10. We then computed principal component analysis, neighborhood graph and clustered the data using the Leiden algorithm ^26^. The data was visualized using UMAP projection^25^. When performing batch correction to remove the technical artifacts introduced by the technologies, we replaced the neighborhood graph computation with batch balanced KNN (bbknn) ^76^.

**Table 1.** Summary of samples processed in the study.

| Sample type | Technologies | Samples | Cells that pass QC |
| --- | --- | --- | --- |
| GL261 | • Microfluidic snRNA-Seq | 10x nuclei GL261-1 | 3,333 |
|  |  | 10x nuclei GL261-2 | 960 |
|  | • Microfluidic scRNA-Seq | 10x cells GL261-1 | 3,951 |
|  |  | 10x cells GL261-2 | 4,641 |
|  | • FACS-based scRNA-Seq | FACS GL261-1 | 235 |
|  |  | FACS GL261-2 | 545 |
|  |  | FACS GL261-3 | 549 |
| GL261-GSCs | • Microfluidic snRNA-Seq | 10x nuclei GL261-GSCs-1 | 3,198 |
|  |  | 10x nuclei GL261-GSCs-2 | 2,289 |
|  |  | 10x nuclei GL261-GSCs-3 | 1,782 |
|  | • Microfluidic scRNA-Seq | 10x cells GL261-GSCs-1 | 3,402 |
|  |  | 10x cells GL261-GSCs-2 | 5,605 |
|  |  | 10x cells GL261-GSCs-3 | 7,025 |
|  |  | 10x cells GL261-GSCs-4 | 4,030 |
|  | • FACS-based scRNA-Seq | FACS GL261-GSCs-1 | 464 |
|  |  | FACS GL261-GSCs-2 | 213 |
|  |  | FACS GL261-GSCs-3 | 188 |
|  |  | FACS GL261-GSCs-4 | 237 |
| Tissue | • Microfluidic snRNA-Seq | Early control-1 | 4,727 |
|  |  | Early control-2 | 4,451 |
|  |  | Late control-1 | 4,207 |
|  |  | Late control-2 | 3,886 |
|  |  | Early peptide-1 | 3,365 |
|  |  | Early peptide-2 | 5,554 |
|  |  | Early peptide-3 | 4,499 |
|  |  | Late peptide-1 | 1,314 |
|  |  | Early TMZ-1 | 5,154 |
|  |  | Early TMZ-2 | 2,728 |
|  |  | Early TMZ-3 | 3,774 |
|  |  | Late TMZ-1 | 648 |
|  |  | Late TMZ-2 | 1,109 |
|  |  | Late TMZ-3 | 5,156 |

## Data availability

The transcriptomics data are available in the public functional genomics data repository Gene expression Omnibus (GEO) identifier:

GSE246154 - Single-nucleus RNA-sequencing (snRNA-Seq) of brain cells from tumor-bearing C57BL/6 mice- https://www.ncbi.nlm.nih.gov/geo/query/acc.cgi?acc=GSE246154

GSE244301 - Single-cell RNA-sequencing (scRNA-Seq) of GL261 and GL261-GSCs; 10X- https://www.ncbi.nlm.nih.gov/geo/query/acc.cgi?acc=GSE244301

GSE246262 - Single-cell RNA-sequencing (scRNA-Seq) of GL261 and GL261-GSCs; smartSeq- https://www.ncbi.nlm.nih.gov/geo/query/acc.cgi?acc=GSE246262

GSE245263 - Spatial transcriptomics of brains from tumor-bearing C57BL/6 mice- https://www.ncbi.nlm.nih.gov/geo/query/acc.cgi?acc=GSE245263

The code is available at github: https://github.com/laugarvi/GL261-GSC-GBM

Tables S1-S15 contain complete analyses performed in the present study.

## Acknowledgements

This work was funded by the grant FEDER RTI2018-099873-B-I00 (Arantxa Tabernero) funded by MCIN/AEI/ 10.13039/501100011033 and “ERDF A way of making Europe”, by the grant PDC2022-133652-I00 (Arantxa Tabernero) funded by MCIN/AEI/ 10.13039/501100011033 and “European Union NextGenerationEU/PRTR”, Junta de Castilla y León, FEDER SA125P20 (Arantxa Tabernero), and Chan Zuckerberg Biohub SF (Norma Neff). L. García-Vicente was supported by the Spanish Ministerio de Universidades. A. Álvarez-Vázquez and R. Flores-Hernández were fellowship recipients from the Junta de Castilla y León and the European Social Fund. We are grateful to Spyros Darmanis and Ashley Byrne for initial support and guidance to the project. We thank T. del Rey for technical assistance.

## Author contributions

L.G.V. contributed to the experimental design and development, data acquisition, and analysis and interpretation of the data. L.G.V., A.A.V., and R.F.H. performed cell culture and in vivo procedures. L.G.V., M.B., and V.T. performed snRNA-Seq experimental procedures. A.G., A.M., Y-J.K., L.D, and A.P. provided computational pipelines and helped with data analysis. A.D., H.M., and S.P. performed sequencing. N.N. helped to design snRNA-Seq experiments and supervised them. A.T. conceived the study, supervised the experimental development, and interpreted the data. L.G.V. and A.T. drafted the article. All the co-authors revised the article for important intellectual content and approved the final version for publication.

## Competing interests

The authors declare no competing interests.

## Notes

https://www.ncbi.nlm.nih.gov/geo/query/acc.cgi?acc=GSE246154

https://www.ncbi.nlm.nih.gov/geo/query/acc.cgi?acc=GSE244301

https://www.ncbi.nlm.nih.gov/geo/query/acc.cgi?acc=GSE246262

https://www.ncbi.nlm.nih.gov/geo/query/acc.cgi?acc=GSE245263

https://github.com/laugarvi/GL261-GSC-GBM

### Competing Interest Statement

The authors have declared no competing interest.

