## Supplementary material for "Single-nucleus RNA sequencing provides insights into the GL261-GSC syngeneic mouse model of glioblastoma": Suppl. information

##### Supplementary Tables

**Table S1.** Differentially expressed genes (DEGs) per cluster (cell types) shown in Figure 2B.

**Table S2.** Differentially expressed genes (DEGs) tumor vs healthy area obtained by Visium platform (t-test).

**Table S3.** Differentially expressed genes (DEGs) between GL261, GL261-GSCs and implanted GL261-GSCs, as shown in Figure 3B and suppl. Figure 2.

**Table S4.** Gene Set Enrichment Analysis (GSEA) in cultured GL261, shown in Figure 3C.

**Table S5.** Gene Set Enrichment Analysis (GSEA) in cultured GL261-GSCs, shown in Figure 3C.

**Table S6.** Gene Set Enrichment Analysis (GSEA) in implanted GL261-GSCs, shown in Figure 3C.

**Table S7.** Shared genes between GL261, GL261-GSCs and implanted GL261-GSCs, related to Figure 3G.

**Table S8.** Gene Set Enrichment Analysis (GSEA) in intracranially implanted GL261-GSCs vs cultured GL261-GSCs.

**Table S9.** Differentially expressed genes (DEGs) in brain tumor-associated immune cells, 7 vs 28d post GL261-GSCs implantation. Related to Figures 5 D-F and suppl. 5C-D.

**Table S10.** Gene Set Enrichment Analysis (GSEA) in brain tumor-associated immune cells at 7d, as shown in Figure 5I.

**Table S11.** Gene Set Enrichment Analysis (GSEA) in brain tumor-associated immune cells at 28d, as shown in Figure 5I.

**Table S12.** Differentially expressed genes (DEGs) in brain tumor-associated immune cells at 7d, under TMZ and TAT-Cx43 peptide treatments, shown in Figure 7C and Suppl. 6C.

**Table S13.** Gene Set Enrichment Analysis (GSEA) in immune cells from brain tumors at 7d postimplantation treated with peptide TAT-Cx43, shown in Figure 7D.

**Table S14.** Gene Set Enrichment Analysis (GSEA) in immune cells from brain tumors at 7d postimplantation treated with TMZ.

**Table S15.** Differentially expressed genes (DEGs) in brain tumor cells at 28d, control vs TMZ treatment, shown in Figure 7H and Suppl. 6D.

#### Supplementary Figure Legends

**Suppl. Figure 1. Quality control metrics of snRNA-Seq data.**

(A) Violinplots showing number of genes, total number of counts, and number of mitochondrial counts captured per cell for each sample.
(B) UMAP visualization colored by sample.
(C) Number of cells analyzed per cell type

**Suppl. Figure 2. Expression of marker genes in GL261 cells.**

UMAP and violin plots show the expression of selected gene markers corresponding to Figure 3B in GL261, GL261-GSCs and implanted GL261-GSCs.

**Suppl. Figure 3. Comparison of the data obtained with different scRNA-Seq technologies.**

(A) PCA visualization of cells analyzed with Smart-Seq2 and 10x methods.
(B) Percentage of cells in each phase of the cell cycle with different technologies.
(C) Boxplots showing the number of counts and number of genes captured per cell with each technology for GL261 and GL261-GSCs. The number of cells analyzed are indicated.
(D) Correlation plots showing similarity between GL261 (left) and GL261-GSC (right) samples taken with different technologies.

**Suppl. Figure 4. Comparison of upregulated genes and pathways captured with different technologies.**

(A) Venn diagrams showing the overlap between differentially expressed genes for each culture type across the three methods (Smart-Seq2, 10x nuclei, and 10x cells). Differential gene expression between GL261-AC and GL261-NS was analyzed independently for each technology using Wicoxon test with a p-value cutoff=0.05.
(B) Stacked bar plots showing the percentage of genes captured with each technology for the top 30 enriched pathways in GL261 (upper) and GL261-GSC (lower).

**Suppl. Figure 5. Expression of immune cell markers.**

(A) UMAP showing the expression of immune cell type gene markers in the immune cell cluster.
(B) UMAP showing classification of cells into cell cycle phase.
(C) UMAP and violinplots show the expression of selected gene markers corresponding to Figure 5G in immune cells at 7 and 28 days post-implantation.
(D) UMAP showing the expression of indicated genes in the immune cell cluster.

**Suppl. Figure 6. Effect of Tat-Cx43$_{266\text{-}283}$ and temozolomide treatment on immune and tumor cells at 7- and 28-days post-implantation (dpi).**

(A) UMAP showing the expression of immune cell type gene markers in the immune cell cluster.
(B) UMAP showing classification of cells into cell cycle phase.
(C) UMAP and violinplots show the expression of selected gene markers corresponding to Figure 5G in immune cells at 7- and 28-days post-implantation.

#### Suppl. Fig. 1

A

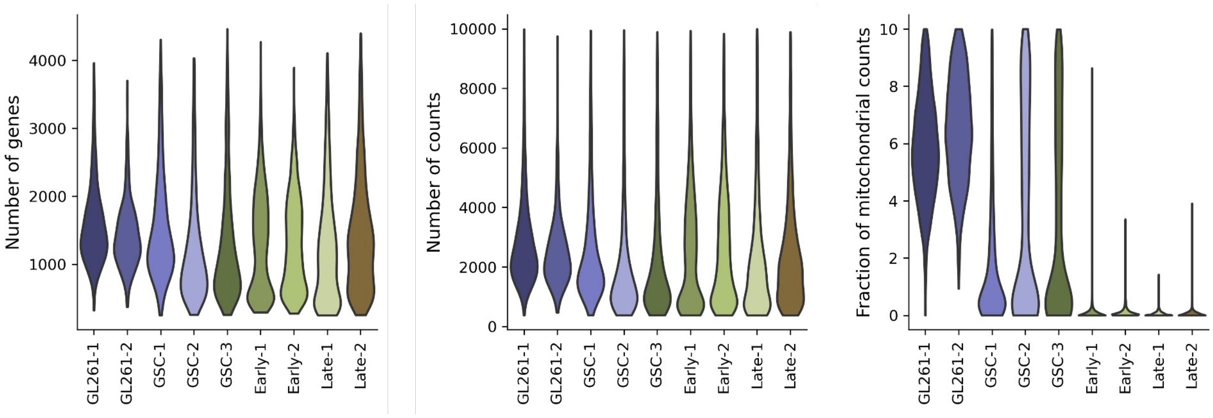


B

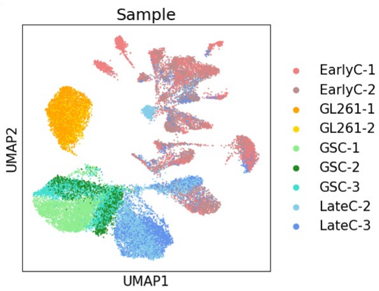


C

| Cell type | Clusters | Number of cells |
|---|---|---|
| GL261-GSCs | 1, 2, 6, 11 | 7268 |
| Neurons | 3, 9, 12, 13, 15, 17, 18, 19, 21, 22 | 6891 |
| Implanted GL261-GSCs | 0, 19 | 5196 |
| GL261 | 4, 5 | 4293 |
| Immune | 7, 10 | 2427 |
| Oligodendrocytes | 8 | 1393 |
| Astrocytes | 14 | 569 |
| OPCs | 16 | 435 |
| Endothelial | 20 | 361 |

Suppl. Fig. 2

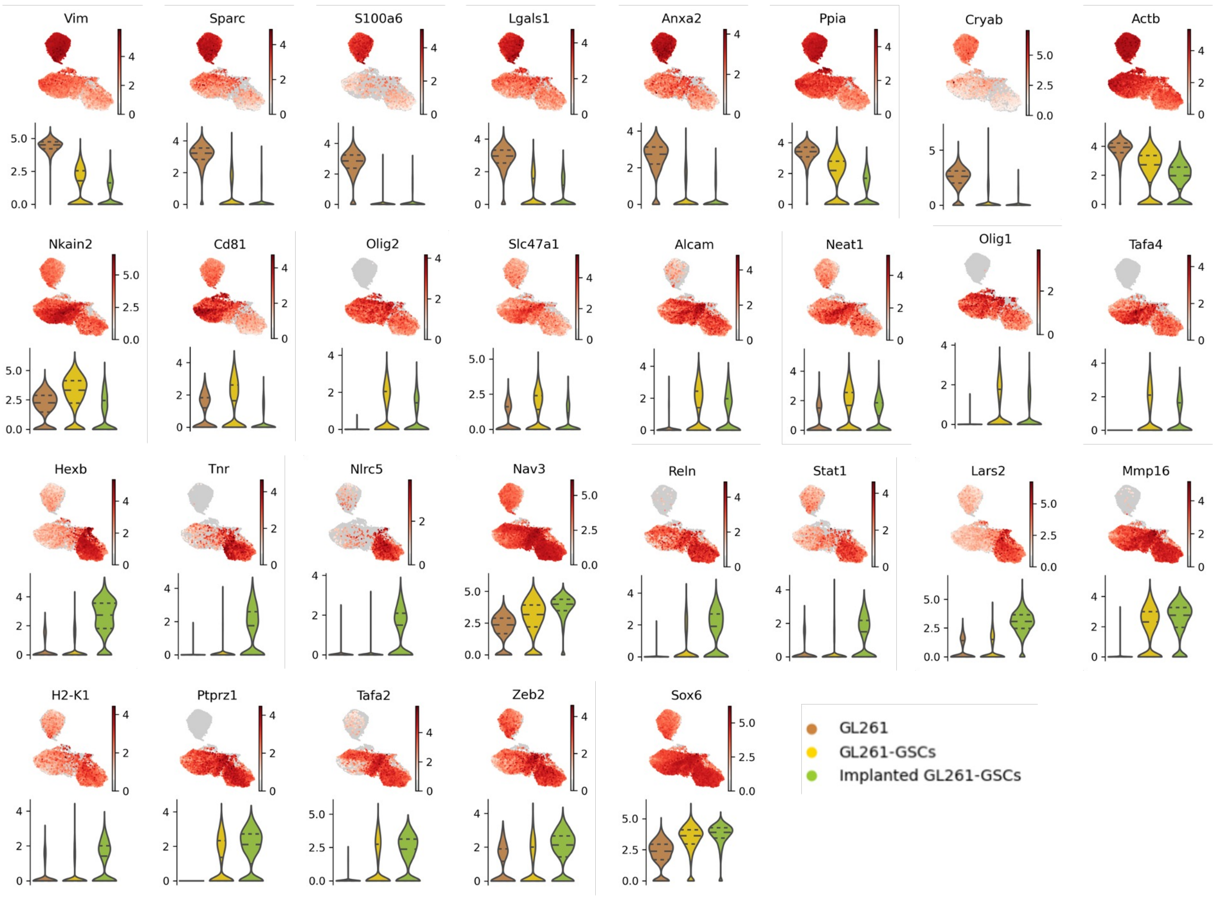
Vim
Sparc
S100a6
Lgals1
Anxa2
Ppia
Cryab
Actb
Nkain2
Cd81
Olig2
Slc47a1
Alcam
Neat1
Olig1
Tafa4
Hexb
Tnr
Nlrc5
Nav3
Reln
Stat1
Lars2
Mmp16
H2-K1
Ptprz1
Tafa2
Zeb2
Sox6
GL261
GL261-GSCs
Implanted GL261-GSCs

Suppl. Fig. 3

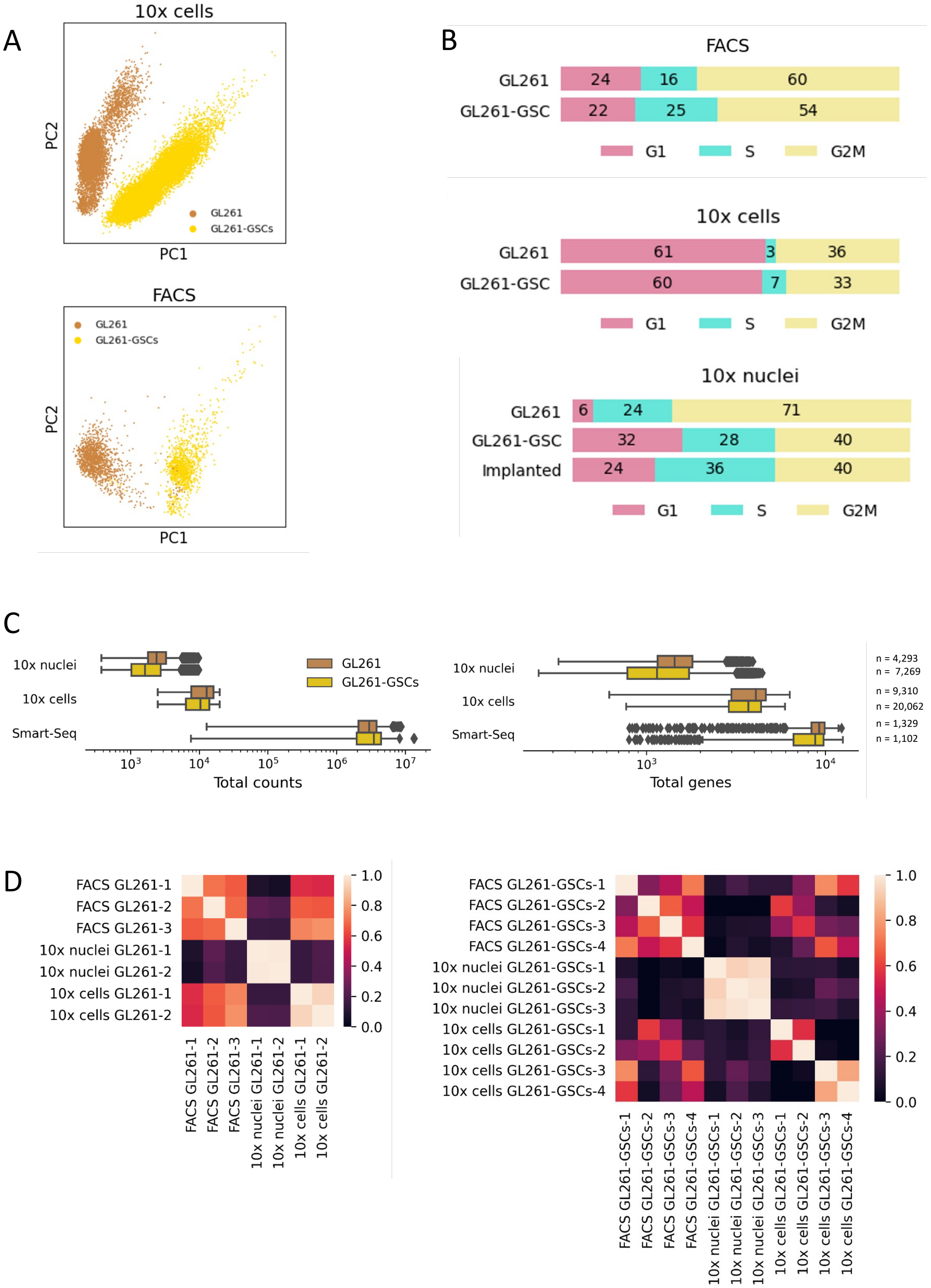
A
10x cells
PC2
PC1
GL261
GL261-GSCs
FACS
PC2
PC1
GL261
GL261-GSCs
B
FACS
GL261 24 16 60
GL261-GSC 22 25 54
G1 S G2M
10x cells
GL261 61 3 36
GL261-GSC 60 7 33
G1 S G2M
10x nuclei
GL261 6 24 71
GL261-GSC 32 28 40
Implanted 24 36 40
G1 S G2M
C
10x nuclei
10x cells
Smart-Seq
GL261
GL261-GSCs
Total counts
10x nuclei
10x cells
Smart-Seq
Total genes
n = 4,293
n = 7,269
n = 9,310
n = 20,062
n = 1,329
n = 1,102
D
FACS GL261-1
FACS GL261-2
FACS GL261-3
10x nuclei GL261-1
10x nuclei GL261-2
10x cells GL261-1
10x cells GL261-2
FACS GL261-GSCs-1
FACS GL261-GSCs-2
FACS GL261-GSCs-3
FACS GL261-GSCs-4
10x nuclei GL261-GSCs-1
10x nuclei GL261-GSCs-2
10x nuclei GL261-GSCs-3
10x cells GL261-GSCs-1
10x cells GL261-GSCs-2
10x cells GL261-GSCs-3
10x cells GL261-GSCs-4

Suppl. Fig. 4

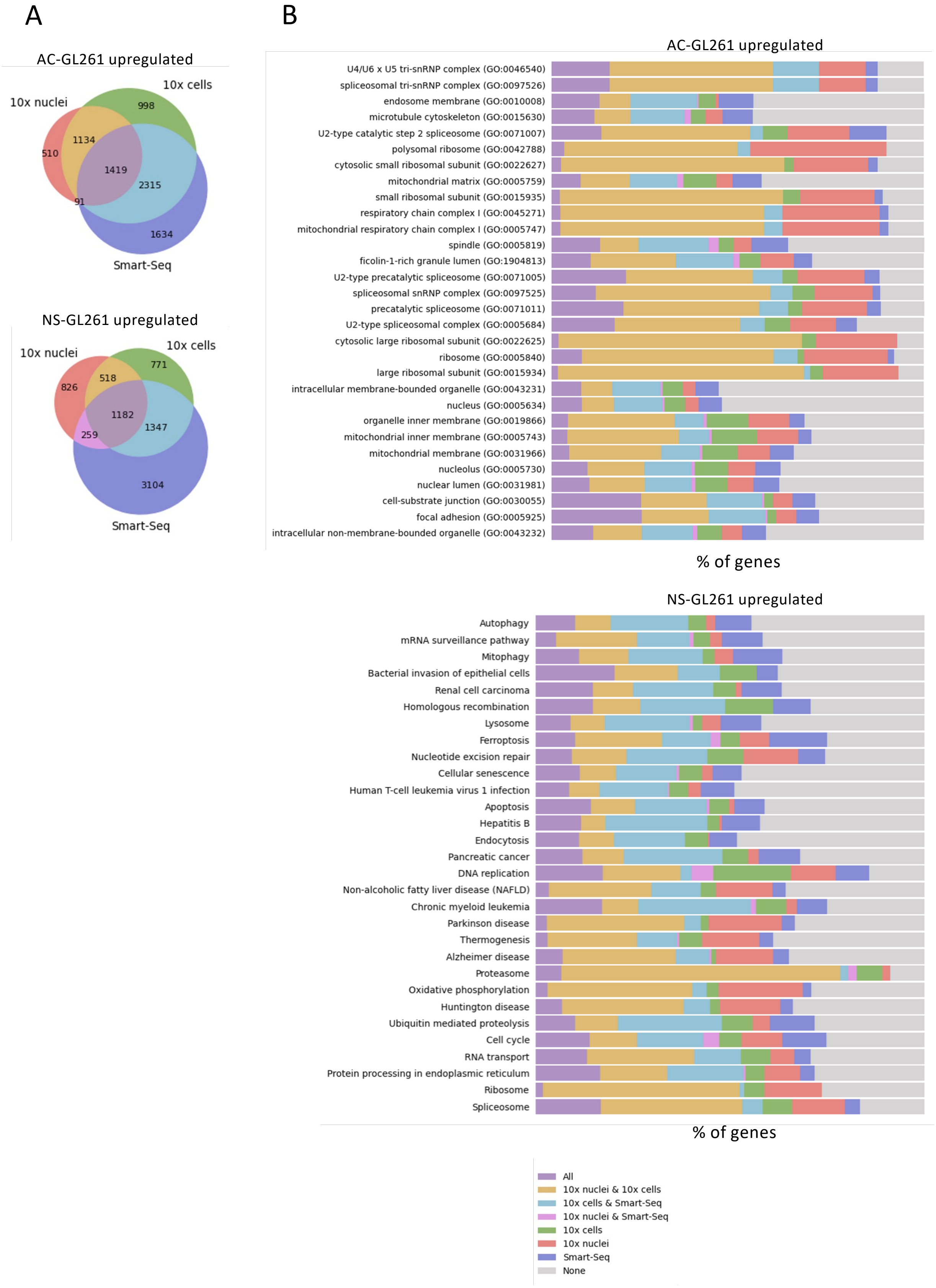
A
AC-GL261 upregulated
10x nuclei
10x cells
Smart-Seq
510
1134
998
1419
2315
91
1634
NS-GL261 upregulated
10x nuclei
10x cells
Smart-Seq
826
518
771
1182
259
1347
3104
B
AC-GL261 upregulated
U4/U6 x U5 tri-snRNP complex (GO:0046540)
spliceosomal tri-snRNP complex (GO:0097526)
endosome membrane (GO:0010008)
microtubule cytoskeleton (GO:0015630)
U2-type catalytic step 2 spliceosome (GO:0071007)
polysomal ribosome (GO:0042788)
cytosolic small ribosomal subunit (GO:0022627)
mitochondrial matrix (GO:0005759)
small ribosomal subunit (GO:0015935)
respiratory chain complex I (GO:0045271)
mitochondrial respiratory chain complex I (GO:0005747)
spindle (GO:0005819)
ficolin-1-rich granule lumen (GO:1904813)
U2-type precatalytic spliceosome (GO:0071005)
spliceosomal snRNP complex (GO:0097525)
precatalytic spliceosome (GO:0071011)
U2-type spliceosomal complex (GO:0005684)
cytosolic large ribosomal subunit (GO:0022625)
ribosome (GO:0005840)
large ribosomal subunit (GO:0015934)
intracellular membrane-bounded organelle (GO:0043231)
nucleus (GO:0005634)
organelle inner membrane (GO:0019866)
mitochondrial inner membrane (GO:0005743)
mitochondrial membrane (GO:0031966)
nucleolus (GO:0005730)
nuclear lumen (GO:0031981)
cell-substrate junction (GO:0030055)
focal adhesion (GO:0005925)
intracellular non-membrane-bounded organelle (GO:0043232)
% of genes
NS-GL261 upregulated
Autophagy
mRNA surveillance pathway
Mitophagy
Bacterial invasion of epithelial cells
Renal cell carcinoma
Homologous recombination
Lysosome
Ferroptosis
Nucleotide excision repair
Cellular senescence
Human T-cell leukemia virus 1 infection
Apoptosis
Hepatitis B
Endocytosis
Pancreatic cancer
DNA replication
Non-alcoholic fatty liver disease (NAFLD)
Chronic myeloid leukemia
Parkinson disease
Thermogenesis
Alzheimer disease
Proteasome
Oxidative phosphorylation
Huntington disease
Ubiquitin mediated proteolysis
Cell cycle
RNA transport
Protein processing in endoplasmic reticulum
Ribosome
Spliceosome
% of genes
All
10x nuclei & 10x cells
10x cells & Smart-Seq
10x nuclei & Smart-Seq
10x cells
10x nuclei
Smart-Seq
None

Suppl. Fig. 5

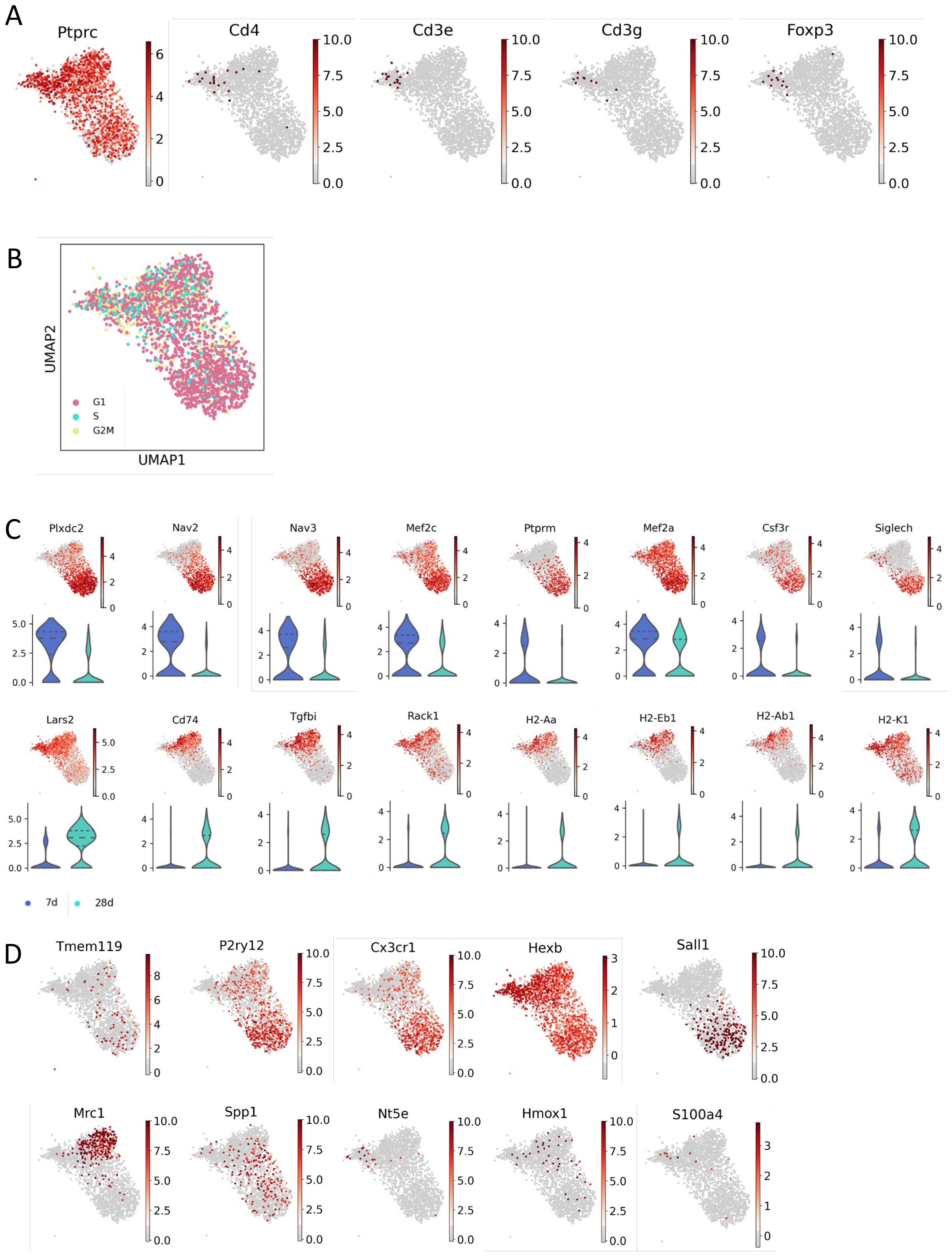
A
Ptprc
Cd4
Cd3e
Cd3g
Foxp3
B
UMAP2
UMAP1
G1
S
G2M
C
Plxdc2
Nav2
Nav3
Mef2c
Ptprm
Mef2a
Csf3r
Siglech
Lars2
Cd74
Tgfbi
Rack1
H2-Aa
H2-Eb1
H2-Ab1
H2-K1
7d
28d
D
Tmem119
P2ry12
Cx3cr1
Hexb
Sall1
Mrc1
Spp1
Nt5e
Hmox1
S100a4

### Suppl. Fig. 6

A

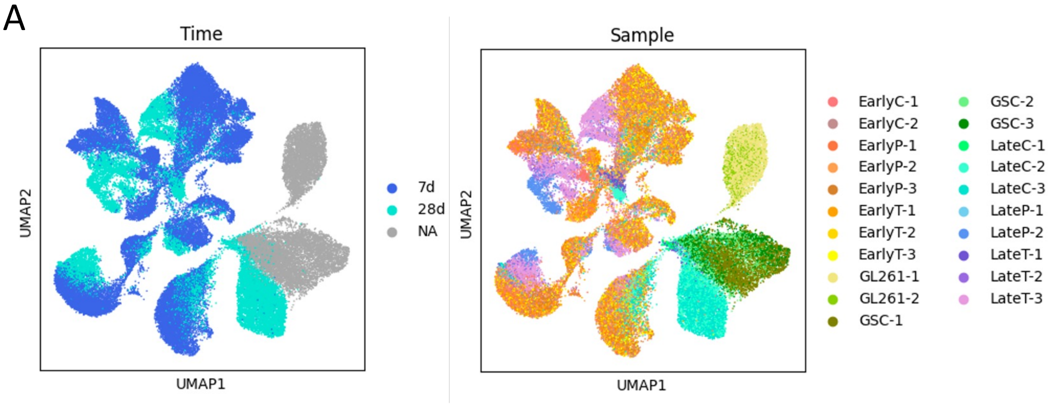


B

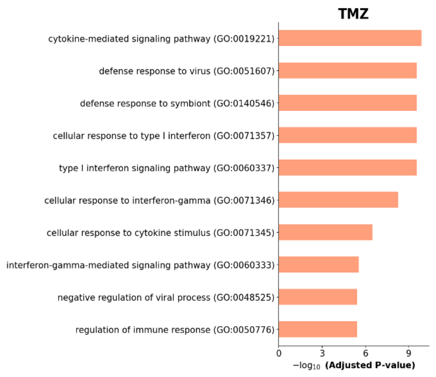


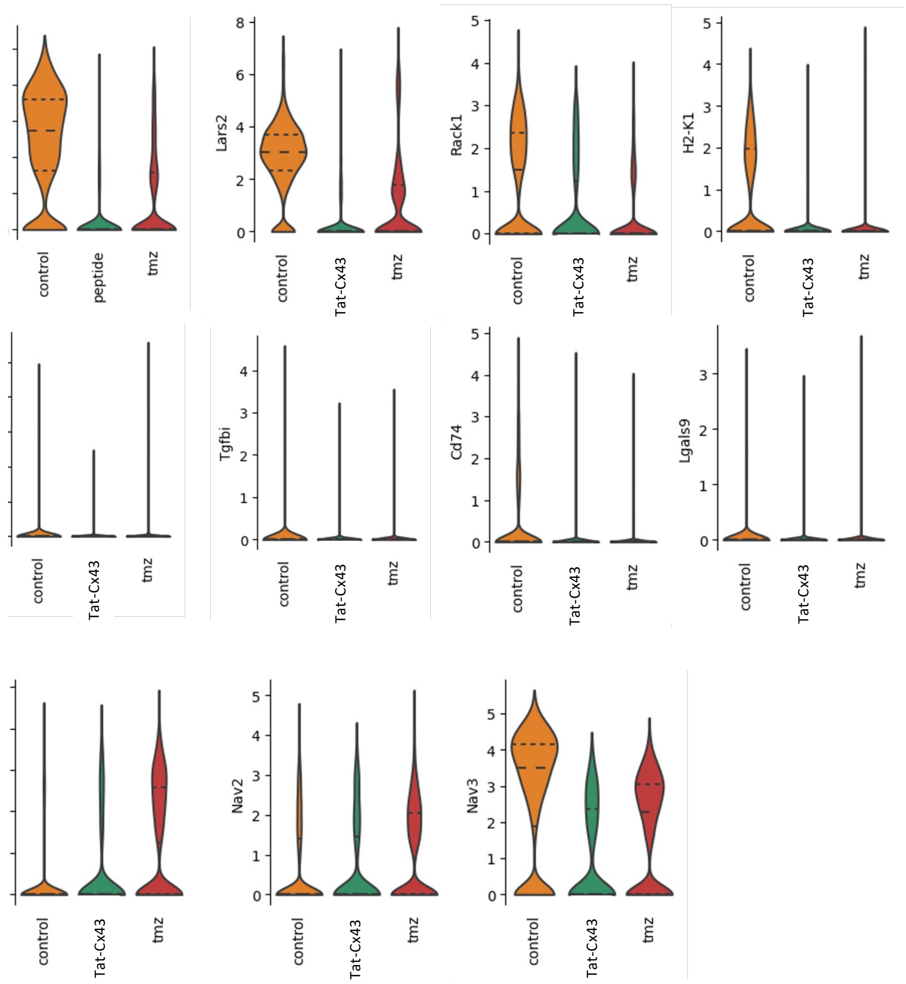


D

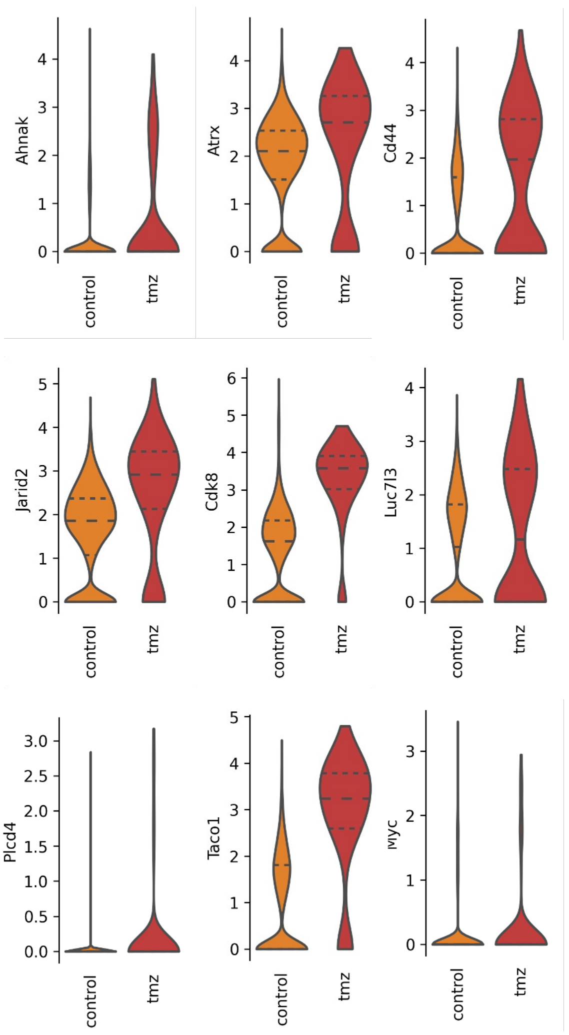
